# CD4+ T-cells create a stable mechanical environment for force-sensitive TCR:pMHC interactions

**DOI:** 10.1101/2024.12.18.629139

**Authors:** Lukas Schrangl, Florian Kellner, René Platzer, Vanessa Mühlgrabner, Paul Hubinger, Josephine Wieland, Reinhard Obst, José L. Toca-Herrera, Johannes B. Huppa, Gerhard J. Schütz, Janett Göhring

## Abstract

Mechanical forces acting on ligand-engaged T-cell receptors (TCRs) have previously been implicated in T-cell antigen recognition and ligand discrimination, yet their magnitude, frequency, and impact remain unclear. We quantitatively assess forces across various TCR:pMHC pairs with different bond lifetimes at single-molecule resolution, both before and during T-cell activation, on platforms that either include or exclude tangential force registration. Our results imply that CD4+ T-cell TCRs experience significantly lower forces than previously estimated, with only a small fraction of ligand-engaged TCRs being subjected to these forces during antigen scanning. These rare and minute mechanical forces do not impact the global lifetime distribution of the TCR:ligand bond. We propose that the immunological synapse is created as biophysically stable environment to prevent pulling forces from disturbing antigen recognition.

## Main Text

### Introduction

The adaptive leg of immune surveillance is realized by the interplay between antigen-presenting cells (APCs) and lymphocytes. Once primed in secondary lymphoid organs, T-cells become highly motile to patrol tissues in search of pathogen-derived antigens presented on the cell surface of APCs by major histocompatibility complexes (MHCs)(*1*). When measured *in vitro*, T-cell receptor (TCR) interactions with nominal peptide/MHCs (pMHCs) are typically of moderate affinity. Yet, T-cells can detect the presence of even a single antigenic pMHC among thousands of non-stimulatory but structurally similar ligands (*2*, *3*). Upon encountering antigen, T-cells rapidly initiate via their triggered TCRs intracellular signaling cascades and establish through a complex interplay of TCRs, accessory molecules and the underlying cytoskeleton an elaborate bi-membrane T-cell:APC interface, termed the immunological synapse (*4*). Its unique properties and temporal plasticity are likely to influence the dynamics of receptor-ligand interactions (*5*). For example, massive cytoskeletal rearrangements (*5*) after initial contact eventually result in the molecular segregation of membrane receptors and ligands based on the size of their extracellular domains (*6*). Furthermore, during activation TCRs are transported from the periphery to the center of the synapse (*7*, *8*), potentially straining the TCR:pMHC bond (*9–11*).

While intrinsic properties of the receptor-ligand bond appear only in part to determine the signaling outcome, biophysical parameters such as mechanical forces have also been implicated (*12–19*). Exerting defined mechanical strain onto the TCR-pMHC bond revealed that molecular force indeed can activate T-cells (*13*, *14*) and even improve ligand discrimination (*20–25*) by increasing (catch bond) or decreasing (slip bond) binding lifetimes (*26–28*). Catch bond behavior was reported with a maximum lifetime increase at 10-15 pN per bond for CD8+ (*20*, *21*, *23*, *24*, *29*) and 15-17 pN per bond for CD4+ T-cells (*25*). Another study involving MHC class II molecules as ligands for CD8+ T-cells implied dynamic catch bonding for the bimolecular complex (*22*). TCRs of CD8+ T-cells experience a force range of 12-19 pN (*30*) or >4.7 pN (*31*) per bond within the immunological synapse as quantified using DNA-based molecular force sensors. In contrast, the quantification of molecular forces exerted by the TCR itself, revealed that CD4+ T-cells do not necessarily reach the peak force regime of the dynamic catch bond (*11*, *32*). Furthermore, when assessing TCR:pMHC interactions in a cell-free context to preclude contextual accessory interactions, catch bond formation was no longer observed (*33*). Taken together, the physiological significance of mechanical forces exerted and experienced by pMHC-engaged TCRs remains unclear: molecular strain could be a supportive or perturbing factor or even both in T-cell antigen recognition (*34*). The principle of “force-shielding”, i.e. the protection of TCR:ligand pairs from molecular strain by surrounding adhesion molecules such as CD2 and LFA-1 has been suggested in order to create a stable biophysical environment (*34*, *35*).

Here we quantified mechanical forces as they were exerted by single pMHC-engaged TCRs within the immunological synapse of CD4+ T-cells prior to and during T-cell activation. We next correlated observed forces with synaptic lifetimes of the respective TCR.pMHC pairs to assess the possibility of catch–slip bond formation. To this end, we employed single-molecule-sensitive microscopy modalities monitoring the optical parameters of (i) a peptide-based molecular force sensor attached to the base of the ligand of interest and anchored to a surrogate APC, or (ii) a proximity-sensitive sensor to measure lifetimes of receptor-ligand interactions. In this fashion we succeeded in determining TCR-exerted molecular forces as well as their frequency of occurrence in synapses of CD4+ T-cells, their impact on TCR:pMHC lifetimes, and ultimately their relevance for T-cell antigen recognition and activation.

### Results

#### Quantification of TCR-imposed molecular forces with a Molecular Force Sensor platform

To quantitate molecular TCR-imposed forces as they occur within the immunological synapse, we built a molecular force sensor (MFS) platform as previously described (*11*, *36*) (see fig. 1 for a schematic overview). In short, glass-supported lipid bilayers (SLBs) were decorated with murine ICAM-1 and B7- 1, as well as monovalent streptavidin serving as an anchor unit for biotinylated MFSs. The MFS platform supports direct visualization via total internal reflection fluorescence (TIRF) microscopy and quantification of TCR-exerted forces within the single-digit piconewton (pN) range via single-molecule FRET. Two fluorophores constituting a FRET pair were attached to the peptide backbone derived from the flagelliform spider silk protein. The peptide acted as an entropic spring which collapsed in the absence of force giving rise to high FRET efficiency, while force application increased the inter-dye distance resulting in reduced FRET yield. The distal part of the peptide was attached to the base of a TCR ligand such as the MHC class II molecule IE^k^ loaded with moth cytochrome C (MCC) or a recently described high affinity derivate of MCC (<u>a</u>ffinity <u>e</u>nhanced <u>p</u>eptide, AEP) (*37*). Both functional units represent stimulating antigens for 5c.c7 and AND TCR-transgenic CD4+ T-cells, which were investigated in this study. Importantly, the AND TCR possesses a higher affinity for IE^k^/MCC than 5c.c7 TCR (*38*). Sensor integrity, mobility, the stoichiometry of the platform, and the usability of these constructs for single-molecule force analysis have been shown in detail before (*11*) or have been determined accordingly (fig. S1).

**Fig. 1.**
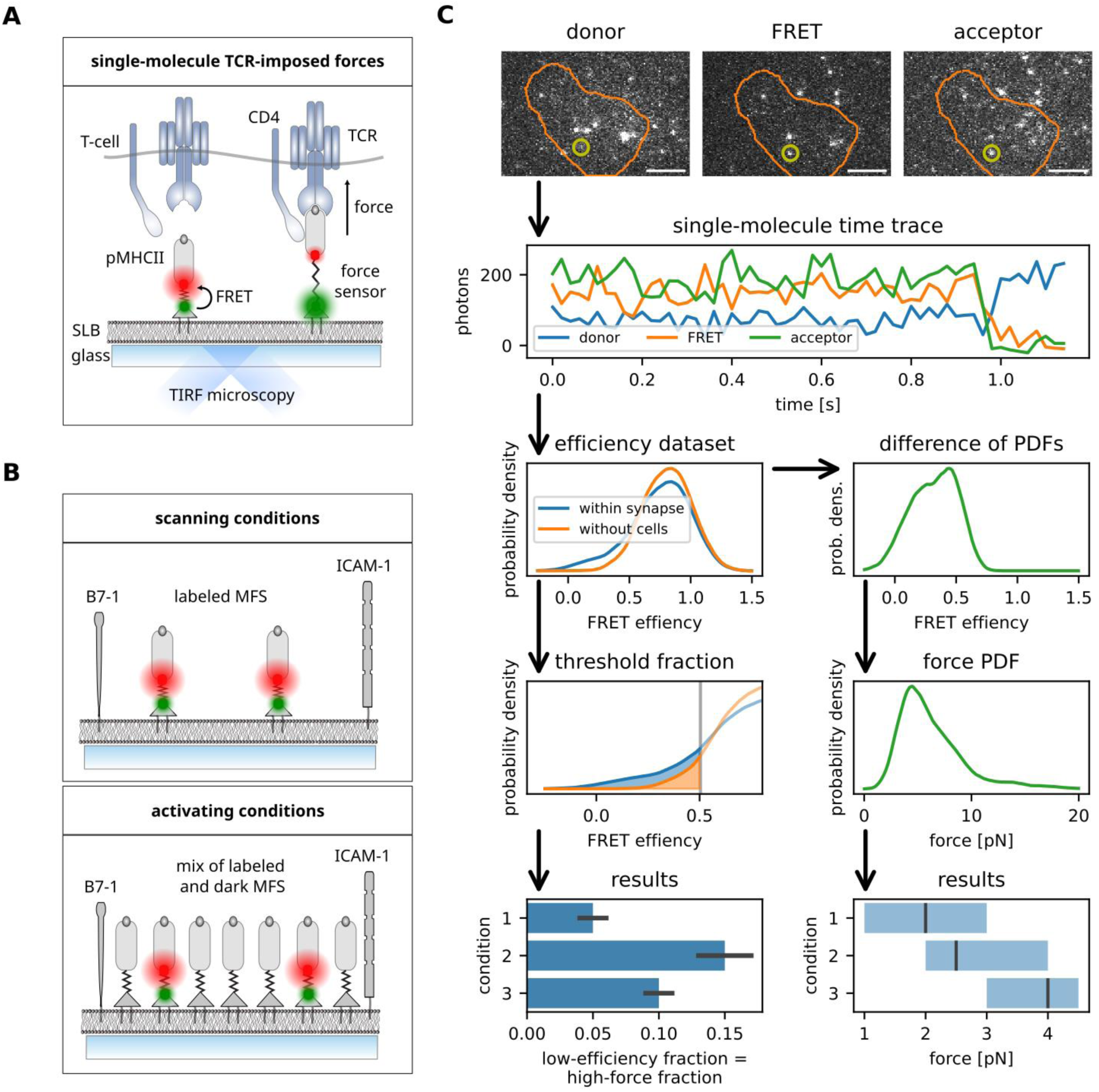
Single-molecule force measurements to quantify TCR-imposed mechanical forces. **(A)** Schematic representation of analog peptide-based molecular force sensor (MFS) platform. His-tagged monovalent streptavidin is directly incorporated into Ni-NTA-DGS-doped supported lipid bilayer (SLB) which can either be fluid or in a gel phase. MFS-conjugated pMHC of interest is anchored via biotin-streptavidin interaction. Single-molecule microscopy (total internal reflection) allows for the measurement of the distance of two fluorophores attached to the peptide backbone of the MFS via Förster resonance energy transfer (FRET). The SLB is also decorated with adhesion (ICAM-1) and costimulatory (B7-1) molecules to promote T-cell adhesion and activation. TCR transgenic T-cells interact with the MFS-coupled pMHC and exert molecular forces extending the spring element of the MFS and increasing the distance between the attached FRET fluorophores, which leads to a decrease in FRET efficiency. **(B)** Schematic representation of activating and scanning conditions. SLBs used for scanning conditions are decorated with fluorescence-labelled MFS at low density, whereas stimulatory SLBs are additionally decorated with non-fluorescent MFS. **(C)** Schematic representation of the data analysis pipeline from data acquisition of single-molecule FRET trajectories to force histograms and statistical data handling. Single-molecule FRET trajectories are transformed into a FRET efficiency histogram. Events with a FRET efficiency below the threshold defined via a 5% false positive rate in data in the absence of cells are counted to compute the proportion of high-force events (plotted as bar graphs, left column). For quantification of single molecule force, the sample and no-cell PDFs are subtracted, and the resulting distribution of the low FRET events are converted into force and displayed as boxplot (right column).

The SLB-based system (fig. 1A) allowed for modulation of lateral ligand mobility by varying the lipid composition giving rise to gel-phase or fluid SLBs, and also the strength of the provided antigenic stimulus determining the level of T-cell activation. Gel-phase SLBs allowed for the registration of tangential and perpendicular pulling forces, whereas the low drag of fluid SLBs permits perpendicular forces only (*11*). SLBs equipped with MFSs at low density (fewer than 0.1/μm^2^), suitable for single-molecule experiments, prompted T-cells to scan for antigen without activation (therefore referred to as “scanning conditions”), while the addition of high-density unlabeled constructs (MFS_0_-IE^k^/MCC) stimulated the T-cells to establish immunological synapses (hence termed “activating conditions”; fig. 1B).

Single-molecule time traces of FRET donor and acceptor molecules were recorded employing alternating laser excitation (*39*) in total internal reflection fluorescence (TIRF) microscopy and subsequently filtered and analyzed (*11*, *40*). To determine the proportion of sensors experiencing forces (“high-force fraction”), first a threshold was derived from data recorded in the absence of T-cells (and hence without force). Datapoints with FRET efficiency below this threshold (note that low efficiency corresponds to high force) were counted towards the high-force fraction (see fig. 1C, left column; details in the methods section). For quantification of the sensors subjected to forces, we estimated their force probability density functions (PDFs) as follows: The FRET efficiency PDF from sensors recorded in the absence of T-cells was scaled and subtracted from the efficiency PDF from sensors imaged within areas of T-cell:SLB contact (Fig. S2, see the methods section for details). The resulting efficiency PDF was transformed using the sensors’ calibration parameters to a PDF of forces (fig. 1C, right column). From this, summary statistics, in particular median and inter-quartile range, were extracted.

#### Binding affinity does not correlate with TCR-imposed mechanical forces

We chose well-characterized TCR:pMHC pairs of different affinity in order to investigate the impact of TCR-exerted strain on bond lifetime. AND and 5c.c7 TCR-transgenic CD4+ T-cells were confronted with IE^k^/MCC-functionalized MFS. Of note, the AND TCR had been reported to exhibit a three times higher 2D affinity towards IE^k^/MCC compared to the 5c.c7 TCR (*38*). We also monitored forces resulting from the interaction between 5c.c7 TCRs and IE^k^/AEP which, based on surface plasmon resonance recordings, feature a 20-fold increased 3D affinity compared to 5c.c7:IE^k^/MCC (*37*). The structural integrity of the newly synthesized MFS-IE^k^/AEP sensor was characterized via size exclusion chromatography (fig. S1A-D). Its ability to efficiently bind and activate T-cells was confirmed via calcium flux analysis (fig. S1E). T-cell’s ability for ligand discrimination was verified for 37°C and room temperature (fig. S1F).

Forces were quantified both for scanning and activating conditions (see fig. 2A–B for a summary of results, figs. S2 and S4 for detailed plots of distributions, and table S1 for summary statistics). Comparing TCR-imposed forces we observe that AND TCRs exerted larger pulling forces than the 5c.c7 TCRs regardless of experimental settings (fig. 2A). For 5c.c7 cells which had been confronted with gel-phase SLBs under scanning conditions or alternatively with fluid-phase SLBs under activating conditions, FRET-based force measurements within synapses or in the absence of T-cells did not give rise to any significant differences (adapted KS-test (*41*), see the methods section), and thus we abstained from further analysis. Surprisingly, the high-affinity 5c.c7:IE^k^/AEP pair experienced even smaller forces than the low-affinity 5c.c7:IE^k^/MCC pair. Only under activating conditions on gel-phase SLBs we observed statistically significant deviations from the baseline. As a control, we performed experiments employing MFSs equipped with H57 TCRβ-reactive single-chain antibody fragments (scF_V_), which exhibit very long lifetimes at room temperature with 50% receptor:ligand bonds still intact after three hours and longer (fig. S3A). Measured forces resembled recordings involving AND T-cells confronted with IE^k^/MCC ligand.

**Fig. 2.**
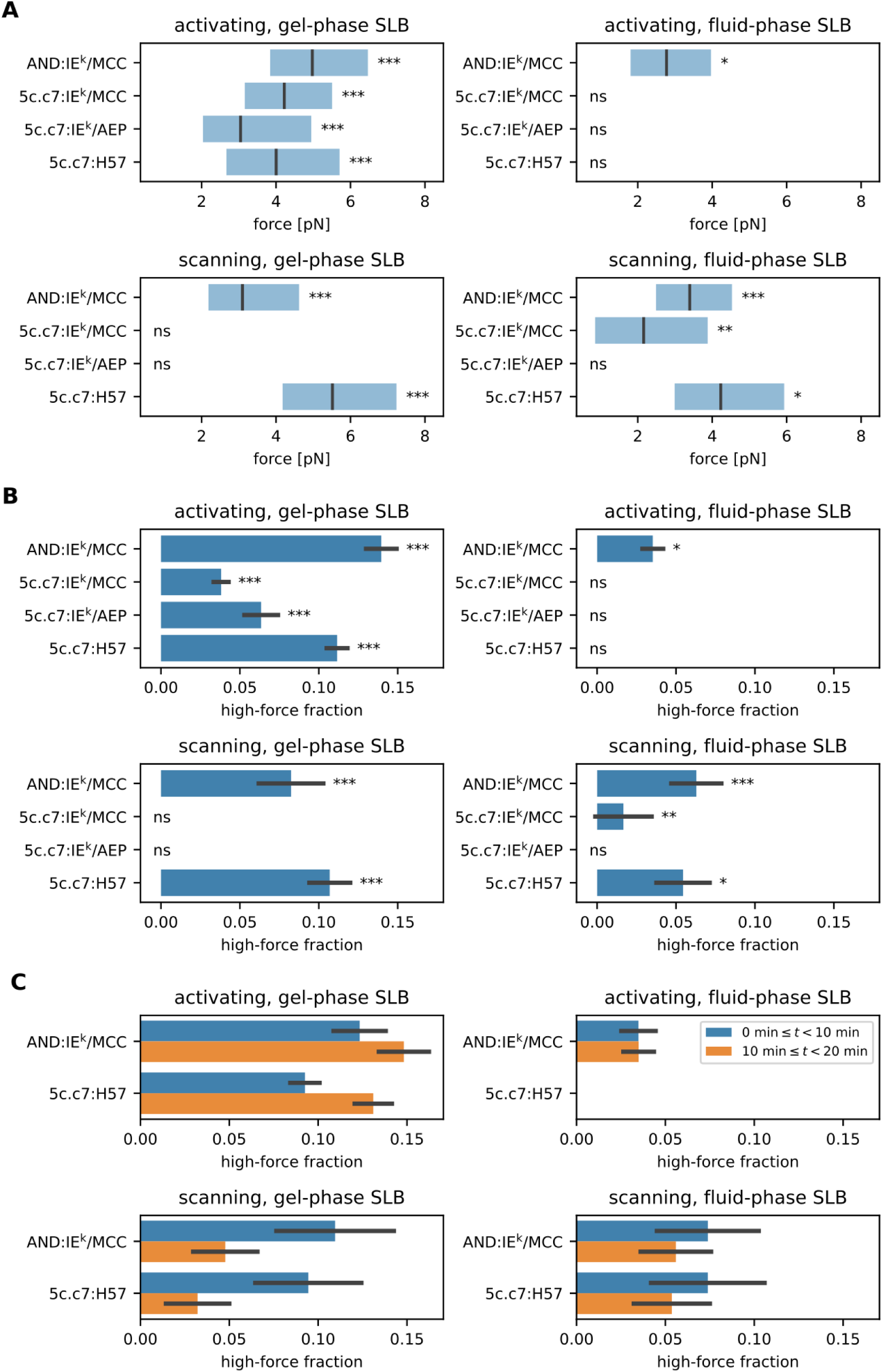
TCR-imposed mechanical forces do not correlate with TCR:pMHC affinity. **(A)** Single-molecule force quantification of investigated TCR:pMHC pairs. The number of trajectories extracted from independent experiments are summarized in Table S1. **(B)** Proportion plot for the recorded single-molecule TCR-imposed forces. All events with FRET efficiency below the calculated threshold were used (see methods). **(C)** Temporal occurrence of the recorded force events in cell seeding intervals (first 10 min after seeding, and 10-20 min after seeding). Significant differences to the no-cell data are indicated by the asterisks in the corner of each plot. * - p-value < 0.05; ** - p-value < 0.01; *** - p- value < 0.001; ns – not significant.

We hence conclude that TCR-exerted mechanical forces within the synaptic environment can reach 6–7 pN (3^rd^ quartile) for the high-affinity AND:IE^k^/MCC and 5c.c7:H57 receptor:ligand pairs, but were substantially reduced if not absent when 5c.c7 T-cells engaged the high affinity ligand IE^k^/AEP or the low affinity version IE^k^/MCC. This may be taken as indication that the interaction of AND TCR with its natural ligand can withstand higher molecular strain than the 5c.c7 TCR.

#### TCR-imposed mechanical pulling forces are rare events

We next examined the frequency of TCR-imposed force events. By determining the proportion of force events above a calculated threshold (see fig. 1C, left column and the methods section), we found marked differences upon varying experimental conditions (see fig. 2B). Of note, results were consistent with above observations: The high-force fraction was largest for AND:IE^k^/MCC and 5c.c7:H57 pairs (about 8–14% on gel-phase SLBs) and considerably reduced for the other experimental conditions (0- 6% on gel-phase SLBs, 0–4% on fluid-phase activating SLBs). For T-cells engaging antigen on fluid-phase SLBs under scanning conditions, force events were registered between 2% (5c.c7:IE^k^/MCC) and 6% (AND: IE^k^/MCC) of all recorded smFRET events. Our analysis furthermore supported comprehensive comparison of different SLB compositions: The majority of high-force events was registered on activating gel-phase SLBs. A similar proportion could be seen under scanning conditions for AND:IE^k^/MCC and 5c.c7:H57 pairs on gel-phase SLBs, but not for 5c.c7:IE^k^/MCC and 5c.c7:IE^k^/AEP. Forces were hardly detectable under activating conditions involving fluid phase SLBs, in line with a previous study of ours involving the use of T-cells engaging SLB-presented MFS (*11*). Force events recorded under activating conditions preferentially occurred 10 min after cell seeding. In contrast, under scanning conditions force events became apparent within the first 10 min of cell seeding (fig. 2C). Of note, temporal analyses of this type are meaningful only to reveal trends of behavior for lack of synchronized T-cell antigen engagement with a sizable number of incoming T-cells also settling during later stages of data acquisition.

Taken together, the large majority of MFSs failed to experience forces. On gel-phase SLBs, the high-force fraction made up less than 14% of all measured single-molecule FRET events. This proportion was further reduced (less than 6%) on fluid-phase SLBs. We hence conclude that only a small fraction of the observed smFRET events experience a mechanical pull.

#### Bound TCR ligands rarely experience mechanical forces

Next, we set out to determine molecular forces which are solely experienced by ligand-bound TCRs. Freely moving ligands embedded within fluid-phase SLBs are markedly reduced in lateral diffusion upon binding to TCRs within the immunological synapse (*42*, *43*). We took advantage of this behavior to discriminate TCR-bound from free MFS-functionalized ligands. To this end, the smallest enclosing circle was computed for each single-molecule track. If its radius was below a calculated threshold (see the methods section for details), it was considered TCR-bound.

As shown in fig. 3A and fig. S5., we observed the highest proportion of bound sensors upon use of stably binding H57-scF_V_-functionalized MFS (about 75% in activating, more than 50% in scanning conditions). For the different TCR:pMHC pairs, 30–60% of MFS were engaging TCRs under activating conditions, compared to 20–30% under scanning conditions.

**Fig. 3.**
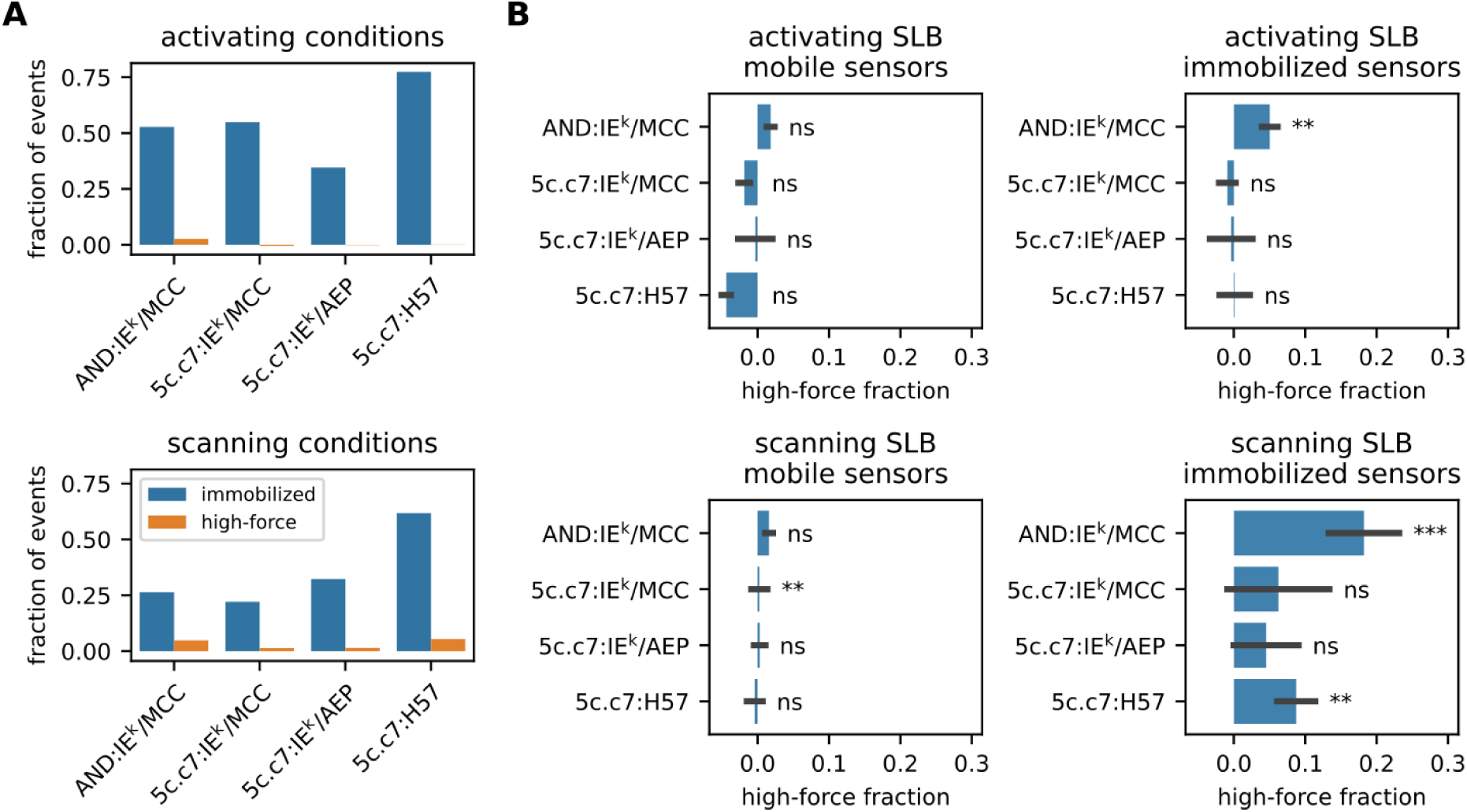
TCR-imposed mechanical forces are rare events. **(A)** Total amount of force events compared to total binding events. The number of trajectories extracted from independent experiments are summarized in Table S1. **(B)** Proportion of the molecular high-force events for the mobile and immobilized fraction of events. Significant differences to the no-cell data are indicated by the asterisks in the corner of each plot. * - p-value < 0.05; ** - p-value < 0.01; *** - p-value < 0.001; ns – not significant.

As expected, only immobilized but not mobile sensors were subjected to forces (fig. 3B). In line with the findings of the previous section, we failed to detect forces under activating conditions, except for the AND TCR for which we noticed a small force-registering fraction. To visualize the number of sensors experiencing forces, we computed the differences of cumulative distribution functions (CDFs) of synaptic sensors and sensors in the absence of cells (fig. S6). For all TCR:ligand pairs measured under activating conditions, CDF differences of bound and unbound sensors appeared visually similar and did indeed not exhibit a significant divergence (adapted KS-test, refer to the Methods section). Under scanning conditions, approximately 20% of bound AND:MFS-IE^k^/MCC and 10% of bound 5c.c7:MFS-H57 events revealed forces (fig. 3B) and CDF differences between mobile and immobilized events were distinct (fig. S6).

In summary, fluid-phase SLBs permitted discrimination between bound and unbound sensors. Under activating conditions, under which T-cells established tight immunological synapses, we failed to detect forces even when restricting the analysis to TCR-engaged sensors. Under scanning conditions, which promoted transient T-cell:SLB contacts, TCR-engaged sensors under force were detectable but more than 1:5 outnumbered by force-free TCR-bound ligands (fig. 3A,B).

#### Global TCR:pMHC interaction lifetime is not affected by TCR-imposed mechanical forces

Finally, we assessed the impact of mechanical forces on the lifetime of synaptic TCR:pMHC interactions. While most protein:protein interactions become destabilized under force (termed slip bonds), stimulatory TCR:pMHC interactions have previously been described as so-called catch bonds, which gain in duration when placed under force (*26–28*). We quantified the average lifetime of TCR:pMHC interactions via single molecule FRET microscopy using fluorescent H57-scF_V_ labeling TCRs and fluorescent SLB-anchored IE^k^/MCC or IE^k^/AEP (*15*, *44*). TCR:pMHC binding leads to measurable FRET by positioning acceptor and donor fluorophores in close proximity (fig. 4A,B). The stability of TCR:H57- scF_V_ bonds was confirmed by varying the incubation temperature (fig. S3A). The binding efficiency of H57-scF_V_ to 5c.c7 and AND transgenic T-cells was tested via flow cytometry and a saturating amount of label was chosen (labeled and unlabeled moieties in a ratio of 1:5, fig. S3B). We measured synaptic TCR:pMHC lifetimes employing activating gel-phase SLBs, i.e. experimental settings giving rise to recordable TCR-imposed forces, and compared them with lifetimes determined using activating fluid phase SLBs, i.e. a regimen for which we failed to detect forces. Of note, this approach is inadequate for measurements carried out under scanning conditions since ligand densities required to record sufficient data points within a feasible time frame are above the T-cell activation threshold (*44*).

**Fig. 4.**
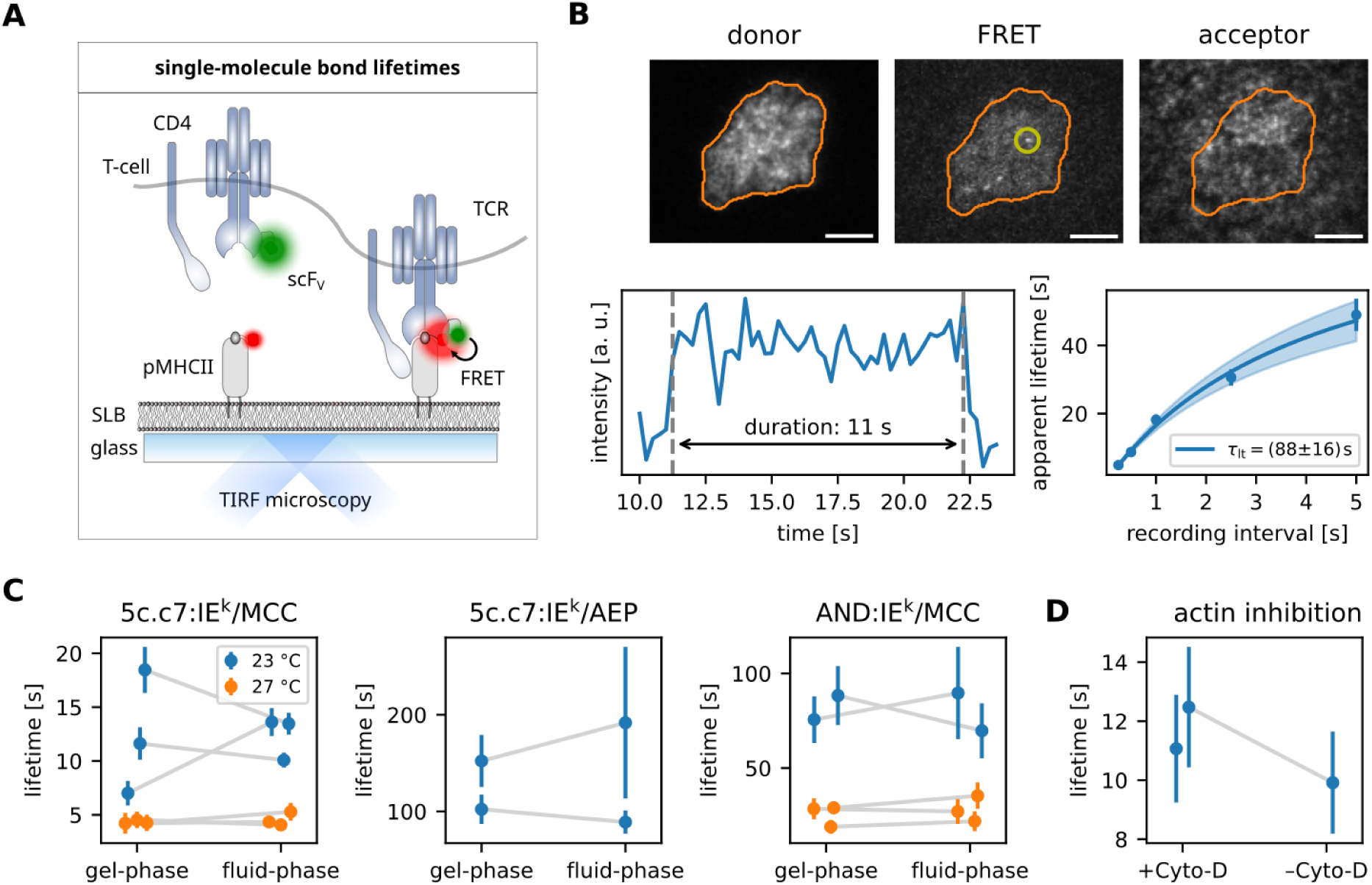
Global TCR:pMHC interaction lifetime is not affected by TCR-imposed mechanical forces. **(A)** Schematic representation of the quantification of single-molecule FRET events for the assessment of TCR:pMHC interaction lifetimes. SLBs were decorated with IE^k^/MCC fluorescently labeled at the C- terminus of the peptide (FRET acceptor). The TCR was labeled via fluorescent H57-scF_V_ (FRET donor). Binding and unbinding could be tracked via the formation of single-molecule FRET signals. **(B)** Representative cell and intensity time trace of a single binding event and its subsequent lifetime analysis. Single molecule FRET events were recorded for different time lags to estimate the effect of photobleaching and to eventually extract the average bond lifetime (*44*). **(C)** Bond lifetime of indicated TCR:pMHC pairs at 23°C and 27°C on gel-phase and fluid SLBs. The results of the individual experimental days are plotted as paired data sets. Mean and standard error of the individual experiments are summarized in table S2. **(D)** 5c.c7:IE^k^/MCC bond-lifetime in the presence and absence of Cytochalasin D (inhibitor of actin polymerization) on gel-phase SLBs at 23°C. Mean and standard error of the individual experiments are summarized in table S3.

Results are summarized in fig. 4C and detailed statistics are shown in table S2. For the 5c.c7:IE^k^/MCC pair, lifetimes were around 10-15 s at 23°C and 5 s at 27°C. Measured lifetimes of synaptic 5c.c7 TCR interactions with IE^k^/AEP amounted to 100 to 200 s (note that for lifetimes this long, the experimental method is inherently imprecise (*44*)). The AND:IE^k^/MCC bond lasted about 75 s at 23°C and 25 s at 27°C. Importantly, lifetimes were not affected by force as the experimental use of fluid-or gel-phase SLBs, which afforded low and high force regimens, respectively, did not change the outcome. In line with this observation, depolymerization of the cortical actin cytoskeleton through cytochalasin D, a measure we chose to eliminate mechanical TCR-imposed forces (*11*), had no noticeable influence on TCR:pMHC lifetime in synapses of T-cells seeded on protein-functionalized gel-phase SLBs.

In summary, we tested various conditions known to evoke different levels of force exertion for their influence on synaptic TCR:pMHC binding kinetics, and found no changes in the global TCR:pMHC bond lifetime, indicating that the rare mechanical force events we observed did not notably impact the overall TCR:pMHC bond lifetime distribution of activating T-cells.

### Discussion

T-cells are highly motile and traverse a multitude of tissues in search of their cognate antigen. It is therefore just a small step to assume that biophysical processes impact antigen scanning efficiency by pushing and pulling on the passing-by T-cells. So far, it has remained an open question whether T-cells exploit TCR-imposed forces for regulatory purposes, or whether they rather stabilize the microenvironment so that receptor-ligand interactions can proceed undisturbed.

In search for an answer, several tools for measuring molecular forces have been developed, which may be categorized according to (i) their reliability to afford single molecule read-out, (ii) analog vs digital read-out, (iii) sensor size, and (iv) the application of external force versus observation of endogenous force. The advantages and limitations of the various methods to quantify TCR-imposed forces have been extensively discussed elsewhere (*45–47*). Our sensor platform is advantageous with regard to all of these aspects, as it permits analog single-molecule readout of TCR-imposed forces while minimizing size-induced side effects within the immunological synapse. Low sensor densities and strict filtering of recorded signal ensured exclusive analysis of single, isolated sensors. As a consequence, issues such as intra-molecular cross-talk, which have been found problematic with other methods (*48*), are circumvented by design. Our molecular force platform has a coiled sized of 2-3 nm and extended can reach 12 nm, i.e. it fulfils the space limitations of the immunological synapse of 13 nm (*49*) for most of the unstretched or slightly stretched states. Still, molecular size has an influence on processes within the immunological synapse and we cannot rule out secondary effects (e.g., on intrinsic force exertion) caused by the insertion of our force sensor. Also, SLBs fail to emulate certain properties of APCs such as 3D structure, protein crowding, and elasticity. However, since current microscopy systems do not offer the required sensitivity for single-molecule force measurements in 3D, we believe that the presented approach is offering the most direct and honest force quantification of surveilling T-cells.

We show in this study that TCRs of CD4+ T-cells impose much lower mechanical forces on their ligands than expected (*25*). While the majority of the TCR-ligand pairs never experienced any force in the observation periods, a small fraction is subjected to minute forces (up to 5.5 pN for the 5c.c7 transgenic TCR, up to 6.5 pN for the AND transgenic TCR). This is considerably below the 15-17 pN range for which maximum discriminatory power has been suggested (*25*). We observed highest TCR-imposed forces for ligands arrested on gel-phase SLBs, and much reduced forces for ligands attached to fluid-phase SLBs. Of note, recorded molecular forces exerted on engaged ligands on mobile platforms were effectively detectable only under conditions allowing antigen surveillance (scanning conditions), but not during immune synapse formation and maintenance (activating conditions). Additionally, as demonstrated previously (*11*), inhibition of actin polymerization abrogates force generation. When measuring TCR:pMHC interaction lifetimes within the immunological synapse in these distinct settings, no differences could be detected. These findings suggest that inside the synapse forces are too rare and/or too weak to substantially influence the TCR:pMHC bond and in particular constitute evidence against catch- and slip bond formation as a fundamental mechanism underlying T-cell antigen recognition. Considering the necessity for T-cells to exactly probe the bond lifetime of their TCR:ligand pairs for ligand discrimination, it seems plausible to prevent mechanical pulling forces from disturbing the interaction in order to increase reproducibility.

With sensors attached to fluid-phase membranes, which offer resistance only in perpendicular direction, synaptic forces appeared as extremely rare events detectable only for the high-affinity AND TCR (with ∼5% of bound sensor data points). In contrast and depending on the TCR:ligand match, up to ∼20% of binding events experienced perpendicular force exertion in synapses of scanning T-cells. This result is surprising in view that T-cells experience global pulling and pushing forces in the nN regime (*9*, *50*). This asymmetry suggests that T-cells employ efficient environment-stabilizing mechanisms in order to prevent excessive molecular forces at their TCR:ligand conjugates once the immunological synapse is established. These observations are in line with the hypothesis of force shielding elements within the immunological synapse and during early T-cell activation (*34*). It is conceivable that the shielding is caused by the large number of adhesion factor interactions which share and buffer the globally experienced mechanical load, thereby efficiently reducing the strain on single TCR:ligand interactions. Hence, the molecular forces a TCR:pMHC bond is experiencing differ considerably depending if they occur in first-contact scenarios without further adhesion control, or if they proceed after immune synapse formation in a more stabilized biophysical environment.

Based on the combined evidence presented here, we propose that T-cells primarily aim to prevent mechanical forces from disrupting molecular interactions by forming immune synapses. While it has long been speculated that T-cells could leverage mechanical forces to discriminate between ligands, this appears feasible, if at all, only during the initial receptor interactions at the microvilli tip as it penetrates the target cell’s glycocalyx. However, direct observation for such a phenomenon is missing. In our study, we demonstrate that, following the initial contact of CD4+ T-cells, most TCR bonds experience minimal strain, and thus do not engage in or benefit from dynamic bonding.

### Methods and Materials

#### Materials

HBSS (Hank’s buffered saline solution, Merck KGaA, Germany); 1xPBS (Merck KGaA, Germany); FCS (fetal calf serum, Biowest, France); DPPC (1,2-dipalmitoyl-sn-glycero-3-phosphocholine; Avanti Polar Lipids, Inc., USA); POPC (1-palmitoyl-2-oleoylglycero-3-phosphocholine; Avanti Polar Lipids, Inc., USA); DGS-NTA(Ni) (1,2-dioleoyl-sn-glycero-3-[(N-(5-amino-1-carboxypentyl)iminodiacetic acid)succinyl] (nickel salt); Avanti Polar Lipids, Inc., USA); DMEM (Dulbeccos’s Modified Eagle’s Medium, Thermo Fisher Scientific, USA); BSA (Bovine serum albumin, Thermo Fisher Scientific, USA); T-cell medium: RPMI 1640 (Life technologies, USA) supplemented with 10 % FCS (Biowest, France), 100 U/mL penicillin/streptomycin (Life technologies, USA), 2 mM L-glutamine (Life technologies, USA), 1mM sodium pyruvate (Life technologies, USA), 0.1 mM non-essential amino acids (Lonza, Switzerland) and 50 µM β-mercaptoethanol (Life technologies, USA); Histopaque-1119 (Merck, USA); Alexa Fluor 555 C2 Maleimide (Thermo Fisher Scientific, USA); Alexa Fluor 647 C2 Maleimide (Thermo Fisher Scientific, USA); ICAM-1-10xHis (Sino Biological, China); B7-1-10xHis (Sino Biological, China); Guanidine hydrochloride (Merck KGaA, Germany); Tris base (Merck KGaA, Germany); NaCl (Merck KGaA, Germany); HRV 3C protease (Pierce, Thermo Fisher Scientific, USA); Dibenzylcyclooctyne-maleimide (DBCO, Jena Bioscience, Germany); Tris(2-carboxyethyl)phosphin-hydrochlorid (TCEP, Merck KGaA, Germany)

#### Animal Model

5c.c7 αβ TCR-transgenic mice (Tg(Tcra5CC7,Tcrb5CC7)IWep, PMID: 1328464) bred onto the B10.A background were housed in groups of 2–5 per cage in the pathogen-free facility at the Medical University of Vienna, Austria. Spleens and lymph nodes were harvested from 12–16 weeks old gender-mixed mice.

Spleens of AND-TCR transgenic B10.BR animals (Tg(TcrAND)53Hed, PMID: 2571940) were removed and sent in DMEM/1% BSA to the Medical University of Vienna on ice. The mice were genotyped by polymerase chain reaction or by cytometry and housed in groups of 2-5 animals per cage in the specific pathogen-free Core Facility Animal Models at the Biomedical Center of LMU Munich, Germany.

Mouse breeding and euthanasia were evaluated by the ethics committees of the Medical University of Vienna and approved by the Federal Ministry of Science, Research and Economy, BMWFW (BMWFW-66.009/0378-WF/V/3b/2016). All procedures to isolate lymphocytes and splenocytes from 8–12 weeks old gender-mixed mice were performed in accordance to Austrian law (Federal Ministry for Science and Research, Vienna, Austria), the guidelines of the Federation of Laboratory Animal Science Associations (FELASA), which match those of Animal Research Reporting In Vivo Experiments (ARRIVE), and the guidelines of the ethics committees of the Medical University of Vienna. Breeding and keeping of AND-TCR transgenic mice has been approved by the Government of Upper Bavaria, protocol 55.2-2532.Vet_02-21-4.

#### Mouse T-cells

Antigen-experienced 5c.c7 and AND murine T-cell blasts were obtained as previously described (*51*). In short, 7.5*10^6^/mL spleenocytes were stimulated with 2 µM C18 reverse-phase high-performance liquid chromatography (HPLC)-purified MCC (88–103) peptide (sequence: ANERADLIAYLKQATK, Intavis, Germany) at 37°C in an atmosphere of 5 % CO_2_ in T-cell medium. Dead cells were removed by centrifugation on a cushion of Histopaque-1119 (Merck, USA) on day 6. T-cells were used for experiments on day 7-9 after initial stimulation.

#### Protein expression and refolding

The TCR β-reactive H57 single-chain fragment (scF_V_) (J0, GenBank: MH045460.1) and the fluorescently labelled H57 scF_V_ (J1, GenBank: MH045461.1) were produced as described (*52*, *53*). In short, scF_V_ constructs were expressed in E. coli and inclusion bodies were extracted. H57 scF_V_s were refolded *in vitro*, concentrated, and purified by gel filtration. The monomeric H57 scF_V_ (J1) was conjugated with Alexa Fluor 555 C2 Maleimide (Thermo Fisher Scientific, USA). Protein-to-dye ratios of site-specifically decorated H57 scF_V_-Alexa Fluor 555 were 1.0.

The murine MHC class II molecule IE^k^ α subunits (with a 6x histidine-tag) and the β subunits (with a 6x histidine-tag) were expressed in E. coli as inclusion bodies and refolded *in vitro* with a placeholder peptide (ANERADLIAYL[ANP]QATK) for later exchange with fluorescently labelled peptides as described (*52*). Refolded IE^k^/ANP was purified via nickel–nitrilotriacetic acid (Ni-NTA)-based affinity chromatography followed by S200 and S75 gel filtration. The peptides MCC (ANERADL<u>IAYLKQATK</u>GGSC), T102S (ANERADL<u>IAYLKQASK</u>GGSC), K99R (ANERADL<u>IAYLRQATK</u>GGSC) Null (ANERAEL<u>IAYLTQAAK</u>GGSC) and AEP (ADGV<u>AFLKAATK</u>GGSC) were site-specifically labelled via Alexa Fluor 647 C2 Maleimide (Thermo Fisher Scientific, USA) and purified as described (*52*). After peptide exchange, fluorescently labelled I-Ek/MCC derivatives were purified via S75 gel filtration and stored at −20 °C in1 x PBS and 50% glycerol before use.

Murine recombinant ICAM-1-10xHis and B7-1-10xHis were purchased from Sino Biological (China).

#### Molecular Force Platform synthesis

Various molecular force sensor (MFS) platforms were created as previously described (*11*). The MFS- IE^k^/AEP construct was newly synthesized and quality controlled according to the established protocols described in (1), whereas the MFS-IE^k^/MCC construct is the same batch as the mentioned publication. In principle, as anchor unit we used a his-tagged monovalent streptavidin which can bind the biotinylated spring unit. The force-sensing unit (spring unit) was covalently attached to the base of a recombinant site-specifically-modified IE^k^/MCC moiety (functional unit). The purified molecular force sensor (spring unit coupled to the functional unit) was subsequently added to SLBs decorated with anchor units.

The anchor unit (recombinant mSAv-3xHis6) was produced as described (*11*, *54*). Biotin-binding deficient subunits (“dead”, with a C-terminal His6-tag) and functional subunits (“alive”, with C-terminal 3C protease cleaving site followed by Glu6-tag) were expressed in E. coli (BL-21) using the pET expression system (Novagen). Inclusion bodies were dissolved at 10 mg/mL in 6 M Guanidine hydrochloride, mixed at a ratio of 1 “alive” to 3 “dead” subunits and refolded in 100 volumes of 1xPBS. The salt concentration was reduced by a series of concentration via a 10 kDa cutoff spin concentrator (Amicon) and dilution with 20 mM TRIS pH 7.0. mSAv-3xHis6 was eluted from a MonoQ 5/50 column (Cytiva) at 20 mM TRIS pH 7.0, 240 mM NaCl. The Glu6-tag was removed via digestion overnight with human 3C protease (Pierce). Size exclusion chromatography (Superdex 200, Cytiva) was used to yield the purified anchor unit.

Fluorescently labelled and unlabelled MFS peptides (spring units) were created as described in (*11*). Sequences: MFS peptide (biotin-N-GG<u>C</u>GS(GPGGA)5GG<u>K</u>YGGS-K(ε-N3)), MFS_C_ peptide (biotin-N-GGGGS(GPGGA)5GG<u>K</u>Y<u>C</u>GS-K(ε-N3)), and MFS_0_ peptide (biotin-N-GGGGS (GPGGA)5 GGKYG GS-K(ε-N3)). The peptides were labelled using Alexa Fluor 555 maleimide (Thermo Fisher Scientific) and Alexa Fluor 647 succinimidyl ester (Thermo Fisher Scientific) at the underlined residues.

TCRβ-reactive H57 antibody single chain fragments with a C-terminal unpaired cysteine residue (H57 cys scFV), subsequently labelled with dibenzylcyclooctyne-maleimide (DBCO, Jena Bioscience), were created as described in (*11*). IE^k^ α featuring an additional C-terminal cysteine residue and IE^k^ β were expressed as inclusion bodies and refolded in the presence of MCC 88-103 peptide (with the sequence H-ANERADLIAYLKQATK-OH) or a recently described high affinity derivate of MCC (AEP, with the sequence H-ADGVAFLKAATK-OH) (*37*) for 2 weeks as previously described (*11*). Correctly folded IE^k^ molecules were isolated via an in-house built 14-4-4 antibody column, reduced with 25 µM TCEP followed by S200 gel filtration. Monomeric IE^k^ MCC with unpaired cysteine residue was collected and coupled to DBCO as described in (*11*).

MFS peptides were mixed with DBCO conjugated proteins and 25 Vol % 2 M TRIS pH 8.0 at a molar ratio of at least 5:1 and incubated for 2 hours (room temperature). A monomeric avidin column (Pierce) was used to remove unreacted protein. After elution with biotin, unconjugated spring was separated from MFS conjugates using size exclusion chromatography (Superdex 75 or 200 column, GE Healthcare).

#### Supported Lipid Bilayer Formation

Gel-phase SLBs were created by using DPPC as carrier lipid, whereas fluid SLBs were created using POPC. Dissolved in chloroform, carrier lipids were mixed with 2 mol-% DGS-NTA(Ni) and subsequently dried under a nitrogen stream for 20 min. After resuspension in 1 mL 1xPBS, they were sonicated for 10 min in an ultrasound water bath (USC500TH, VWR, England) at room temperature (fluid SLBs) or 55°C (gel-like SLBs). The resulting small unilamellar vesicle solution was diluted to 125 µM using 1xPBS.

The original cover slip of an eight-well chamber (Nunc Lab-Tek, Thermo Fisher Scientific, USA) was replaced by attaching a plasma-treated (10 min; PDC-002 Plasma Cleaner, Harrick Plasma, USA) microscopy cover slip (MENZEL-Gläser Deckgläser 24×60mm #1.5) using duplicating silicone (Twinsil soft 18, picodent, Germany). The vesicle solution was incubated for 20 min at room temperature (fluid SLBs) or 55°C (gel-like SLBs) to form lipid bilayers. For calcium flux ligand titration experiments, a 16- well chamber (Nunc Lab-Tek, Thermo Fisher Scientific, USA) was used instead. SLBs were subsequently washed with 1xPBS at room temperature to remove excess vesicles.

His-tagged proteins (for activating conditions: 10 ng mSAv-3xHis6, 30 ng ICAM-1, 50 ng B7-1; for scanning conditions: 0.4 ng mSAv-3xHis6, 30 ng ICAM-1) were added in 50µL 1xPBS to each well and incubated for 60 to 75 minutes at room temperature. Afterwards, the bilayer was washed with 1xPBS. SLBs were finally incubated for 20 minutes with MFS variants (for activating conditions with MFS-H57: 10 ng MFS_0_-H57, 30 to 100 pg MFS-H57 for each experimental day adjusted to reach densities suitable for single molecule experiments; for activating conditions with MFS-IE^k^/MCC or MFS-IE^k^/AEP: 15 to 20 ng MFS_0_-IE^k^/MCC, 30 to 100 pg MFS-IE^k^/MCC or MFS-IE^k^/AEP for each experimental day adjusted to reach densities suitable for single molecule experiments; for scanning conditions: 30 to 100 pg MFS for each experimental day adjusted to reach densities suitable for single molecule experiments) in 50 µL 1xPBS per well for binding to the mSAv-3xHis6. Bilayers were washed to remove excess MFS. Immediately before adding cells, the buffer was exchanged for HBSS containing 2% FCS.

For titration experiments, molecular densities were determined by dividing the fluorescence signal per pixel by the single molecule brightness recorded at the same settings, considering the effective pixel width of 160 nm. In single molecule samples, densities were determined by counting the number of molecules and dividing by the area of the field of view.

#### Single Molecule Microscopy Setup

For live cell imaging with single-molecule resolution, we used objective-type total internal reflection (TIR) illumination of fluorophores which was achieved using an objective with high numerical aperture (*α* Plan-FLUAR 100x/1.45 oil, Zeiss, Germany). Fluorophores were excited with 640nm (OBIS 640, Coherent, USA) or 532 nm (LCX-532L with L1C-AOM, Oxxius, France) laser light coupled into an epifluorescence microscopy (Zeiss, Germany). The emission beam was separated from the excitation light via a quad-band dichroic mirror (Di01-R405/488/532/635-25×36, Semrock Inc., USA).

Using a beam splitter device (Optosplit II, Cairn Research, UK) comprising a dichroic mirror (FF640 - FDi01-25×36, Semrock Inc., USA) and bandpass filters (ET570/60m and ET675/50m, Chroma Technology Corp, USA), the fluorescence emission path was split and projected side-by-side onto the chip of an electron multiplying charge-coupled device (EM-CCD) camera (Andor iXon Ultra 897, Andor Technology Ltd, UK). The microscope and peripherals were controlled by using the SDT-control software developed in-house.

In addition, we used a 405 nm laser (iBeam Smart 405-S, Toptica Photonics AG, Germany) to image Fura-2-loaded cells or reflection interference contrast microscopy (IRM) to visualize the cell contours. For the latter, white light emitted by a mercury arc lamp (HBO 100, Zeiss, Germany) was coupled into the emission beam path via a dichroic mirror (DMSP680B, Thorlabs, Inc., USA) and enabled/disable d via an electronic shutter (VS14, Vincent Associates, USA).

#### Single Molecule FRET Measurement for Molecular Force Quantification

Single molecule force FRET experiments were performed as previously described (*11*, *36*). In summary, T cells were seeded onto sensor-functionalized SLBs. Once cells started spreading (typically after 2 to 3 minutes), image sequences were recorded by alternatingly exciting the donor fluorophore using the 532nm laser and the acceptor using the 640nm laser. For a typical video, this process was repeated 300 times with illumination durations of 5 ms and pauses of 5–85 ms between frames to allow for camera read-out and to adjust the recording rate. In addition, cell outlines were recorded at the beginning and end of each video either using 405 nm excitation (if cells were loaded with Fura-2) or by means of IRM.

For flatfield correction, SLBs with high densities of MFS (5–100/μm²) were employed. First, an image upon 640nm laser excitation was recorded, followed by complete photobleaching of the acceptor fluorophores at high laser intensity. Finally, a frame upon 532 nm excitation was recorded.

For image registration, images of fluorescent beads (TetraSpeck, Invitrogen, USA) were recorded upon 523 nm laser excitation.

#### Single Molecule Force Data Analysis

Data were analyzed as previously described (*11*, *40*). In short, our publicly available software (*55*) was used to perform the following tasks: loading of raw data, registration of donor and acceptor emission channels, localization of fluorescent signals, single-molecule tracking, single-molecule intensity determination, nearest-neighbor analysis to discard overlapping signals, flatfield correction, computation of apparent FRET efficiencies and stoichiometries, stepwise bleaching analysis to remove signals from multiple emitters, computation and application of FRET correction factors (bleed-through, direct acceptor excitation, detection and excitation efficiency factors), filtering based on FRET efficiency and stoichiometry, segmentation of images of cells for cell outline determination, saving of resulting single-molecule data.

#### Database management and evaluation criteria

Experimental data was collected over extended time frames. To guarantee reproducibility and consistency a tight control frame work was established which was adhered to during each experimental day. The collected data from the smForce experiment and the simultaneously measured calcium response data was organized on a file server. After completed quantification of the molecular force and calcium flux response, each data set was evaluated to be part of the integrated result database. Evaluation criteria were the functional state of the T-cell population (positive vs negative control), the functional state of the SLBs (activating vs scanning conditions), the mobility of the mobile SLBs, the absence of force events within the no-cell data, and the response of the positive control of the force experiment (MFS-H57 on gel-phase SLBs). In case of the extraction of immobilized signals from the mobile fraction, the absence of immobilized signals within the no-cell data, was included as another evaluation criterium. Consistently treated data was then pooled to arrive at the final quantification of the molecular force for each investigated condition.

#### Hypothesis tests with single-molecule force data

Datapoints from the same single-molecule track may be correlated, precluding the use of regular statistical tests such as the KS test. Instead, we employed permutation tests and ensured that tracks were kept intact during resampling (*41*). As null hypothesis we defined that FRET efficiency distributions originating from sensors on SLBs without cells are indistinguishable from sensors within the interaction area between cells and SLBs, as alternative hypothesis we stated that TCR-engaged sensors experience greater force and therefore reduced efficiency. Hypotheses were tested using the maximum difference of CDFs as a test statistic in the permutation tests (9999 resamples), thereby effectively performing a one-sided KS test on the tracking data (figures 2 and 3, table S1). The difference between distributions of mobile vs. immobilized sensors (figure S6) was determined likewise.

#### Analysis of the high-force (=low-efficiency) fraction

In order to determine the proportion of single-molecule FRET events originating from sensors subjected to force, first a threshold was computed: From data recorded without cells under otherwise same conditions (i.e., SLBs with the same lipid and protein composition), the 0.05 quantile was determined. Using pooled cell-derived single-molecule data, dividing the number of events *n*_*low*_ with a FRET efficiency below this threshold divided by the total number of events *n*_*total*_ thus yielded the proportion of force-affected sensors with an expected 5% false-positive rate. Therefore, the actual proportion was calculated via *n*_*low*_⁄*n*_*total*_ − 0.05. In order find an estimate for the standard error, the calculation was repeated 1000 times with randomly resampled data. Values present in figures 2 and 3 as well as table S1 are mean and standard deviation of this process. Resampling was performed such that individual single-molecule tracks were kept intact as they may contain correlated data.

To quantify the magnitude of forces experienced by sensors, the distribution of forces corresponding to the high-force fraction was isolated as follows: Average shifted FRET efficiency histograms (range: −0.5 to 1.5, number of shifts: 40, number of bins determined by Sturges’s rule, i.e., 1 + ⌈*log*_2_(*n*)⌉) yielding graphs (i.e., pairs of *x*, *y* coordinates) of the probability density functions (PDFs) of cell-free as well as cell-impacted FRET efficiency data. We subsequently introduced two scaling parameters *a*, *b* for the cell-free PDF. *a* was used to scale the *x* coordinate according to 0.87 + *a*(*x* − 0.87), effectively scaling the peak width around the FRET efficiency of 0.87 (which corresponds to zero force). *b* scaled the peak height via *by*. A non-linear least squares fit (*scipy.optimize.least_squares* function (*56*)) was used to find the best approximation of the scaled cell-free PDF to the cell-impacted PDF in the low-force regime (FRET efficiency > 0.75). The thusly scaled cell-free PDF was then subtracted from the cell-impacted PDF, yielding an estimate of the FRET efficiency PDF of the high-force (= low efficiency) fraction. Using the sensor calibration function, this was converted to a force PDF. Since a PDF has to be strictly non-negative, all negative y values were replaced with 0. Finally, the quartiles as shown in figure 2 and table S1 were computed.

#### Analysis of the mobile fraction

The distinction between mobile and immobilized MFS tracks was made as previously described (*11*). The smallest enclosing circle of the track was computed using the *spatial.smallest_enclosing_circle* function from the *sdt-python* Python package (*57*). Its reduced radius was computed by dividing the radius by the square root of the duration of the track, since the squared radius is expected to grow linearly with time assuming free diffusion. Any track with a reduced radius of less than 0.35 μm/s^0.5^ was considered immobilized. To ensure consistency, all tracks containing less than five observations were discarded.

#### Single Molecule FRET Measurement for Quantification of Interaction Lifetimes

Approximately 20% of TCRs on T-cells were labelled using AF555-H57-scF_V_ in order to reduce bleed-through into the FRET acceptor channel (1 part labelled H57-scF_V_ was pre-mixed with 4 parts unlabelled H57-scF_V_ to reach a final labelling mass of 120 ng per 1 Mio T-cells). Labelling was performed on ice for 20 min, excess scF_V_ was removed by washing with cold imaging buffer (HBSS + 2% FCS). Cells were subsequently kept on ice. The buffer in SLB-containing Labtek wells was exchanged with imaging buffer, and 10^5^ T-cells were seeded immediately before measurement.

smFRET events were recorded using 10 ms illumination time. Acceptor fluorophores were excited using a 640 nm laser at the beginning and the end of the recording, whereas donor fluorophores were excited distinct number of times using varying delays. Lifetimes were recorded at 23°C and 27°C for various receptor:ligand pairs.

#### Lifetime Data Analysis

Data analysis was performed as previously described (*44*). In short, we used our publicly available software (*58*) to load the data, perform image registration for donor and acceptor emission channel, correct acceptor emission images for donor bleed-through, localize and track single-molecule signals, apply changepoint detection to ensure single-step appearance and disappearance of analyzed FRET events, and manually verify each single-molecule trajectory. Track lengths were subjected to survival analysis to determine the apparent lifetime for each recording interval. Fitting apparent lifetime vs. recording interval curves with the appropriate model allowed for disentanglement of unbinding and photobleaching contributions, yielding the characteristic bond lifetime.

#### Calcium Flux measurements

The ratiometric dye Fura-2-AM (Life technologies, USA) was used as an intracellular reporter on calcium levels as previously reported (*59*). 5×10^5^ T-cells were incubated with 2µM Fura-2-AM in T-cell medium for 15 min at room temperature and washed with imaging buffer (HBSS with 2% FCS).

For antigen titration experiments, calcium response was recorded at room temperature, as well as 37°C. Temperature control was carried out with a heating unit and a box enclosing the microscope. 1×10^5^ T-cells were seeded onto SLBs decorated with increasing densities of IE^k^/MCC (agonist), IE^k^/T102S (weak agonist) or IE^k^/Null (non-agonist), as well as unlabeled B7-1 and ICAM-1 (100 µm-2). Up to 12 different conditions were recorded simultaneously using 16-well Lab-Tek Chambers (Nunc). A calcium multiplex setup was used to excite Fura-2-AM using a monochromatic light source (Leica) coupled into an inverted Leica DMI4000B microscope which was equipped with a 20x objective (HCX PL Fluotar 20x, NA = 0.5, Leica), the dichroic beamsplitter FU2 (Leica) and the bandpass filter ET525/36 (Leica). Excitation was continuously switched between 340 nm and 387 nm, which was achieved with the use of a fast excitation filter wheel (Leica) containing the excitation bandpass filters 340/26 and 387/11 (Leica). Images were recorded every 15 seconds for a total recording time of 20 minutes. An automated XY stage (Leica) allowed fast changes between different positions (i.e. wells).

T-cell quality was monitored in parallel to each single-molecule FRET experiment for each experimental day. T-cells were seeded onto functionalized SLBs decorated with either unlabeled B7-1 and ICAM-1 (100 µm-2) as negative control, or additionally unlabeled IE^k^/MCC (100 µm-2) as positive control. Each SLB used for smForce experiments was controlled for its ability to elicit calcium response of seeded T-cells (i.e. activating and scanning conditions) for each experimental day. The ratiometric dye Fura-2 was excited with a monochromatic light source (Polychrome V, TILL Photonics, Germany) coupled to a Zeiss Axiovert 200M, equipped with a 10× objective (UPlanFL N 10x, NA = 0.3, Olympus, Japan), a long-pass filter (T400lp, Chroma Technology, USA), an emission filter ET510/80 (Chroma Technology, USA), a 1.6× tube lens, and an EM-CCD camera (Andor iXon 897, Andor, UK). The dye was excited at 340 and 380 nm with illumination times of 75 and 30 ms for a total recording time of 15 min with a rate of 1 image per second. Experiments were carried out at room temperature.

Calcium image analysis was performed using our custom-made MATLAB (Mathworks, Inc., USA) software described in (*11*).

#### Flow cytometry

For cell surface labelling, 5×10^5^ T-cells were labelled with AF555-H57-scF_V_ (250 µg/mL) for 15 min on ice, and washed 2 times in FACS buffer (1x PBS, 1% BSA, 0.02% NaN3). For titration experiments, the following mass of the AF555-H57-scF_V_ was used: 20, 200, 600, 1800, 5400, and 16200 ng per 1 Mio cells. For time curve experiments 1800 ng per 1 Mio T-cells was used. After washing T-cells were placed on ice, room temperature or into a 37°C water bath. Samples were removed from the respective temperatures after the following intervals (0, 5, 10, 30, 60, and 120 min) and immediately analysed. Unlabelled cells were serving as reference and measured as separate sample. Samples were analyzed on the Cytek Aurora (Cytek Biosciences). Data derived from flow cytometry measurements were analyzed with the FlowJo v10 software (BD Biosciences).

## Funding

Austrian Science Fund (FWF) project P32307-B (LS, JG, GS)

Austrian Science Fund (FWF) project P30214-N36 (LS, GJS)

Austrian Science Fund (FWF) project P25775-B2 (FK, JBH)

German Research Council (DFG) project OB 150/7-1 (JW, RO)

Boehringer Ingelheim Fonds (RP)

Vienna Science and Technology Fund (WWTF) LS13-030 (JG, FK, LS, JBH, GJS)

## Author Contributions

conceptualization: JG, GS, and LS;

data curation: JG and LS;

formal analysis: JG and LS;

funding acquisition: JG, JH, and GS;

investigation: JG, and LS;

methodology: FK, JG, JW, LS, PH, RP, RO, VM;

project administration: JG, GS, and JH;

resources: FK, JG, JW, RO, RP, PH, and VM;

software: LS, and JG;

supervision: JG, GS, and JH;

validation: JG and LS;

visualization: LS, and JG;

writing—original draft preparation: JG and LS;

writing—review and editing: GS, JG, JH, JTH, and LS

All authors have read and agreed to the published version of the manuscript.

## Competing interests

Authors declare that they have no competing interests.

## Data and materials availability

All data generated or analyzed during this study are included in this published article (and its supplementary information files). The unprocessed imaging data of this study is available from the corresponding authors upon request.

## Code availability

We provide the Python code for single-molecule FRET analysis (https://github.com/schuetzgroup/fret-analysis, DOI: 10.5281/zenodo.4604567) as well as the underlying Python library (https://github.com/schuetzgroup/sdt-python, DOI: 10.5281/zenodo.4604495). Additionally, we provide the Python code for single-molecule Lifetime analysis (https://github.com/schuetzgroup/smfret-bondtime, DOI: 10.5281/zenodo.12571064).

## Supplementary Materials

Fig. S1 to S6

Table S1 to S3

## Notes

### Competing Interest Statement

The authors have declared no competing interest.

## References

1. L. Dupré, R. Houmadi, C. Tang, J. Rey-Barroso, T Lymphocyte Migration: An Action Movie Starring the Actin and Associated Actors. Frontiers in Immunology 6, 586 (2015).

2. D. J. Irvine, M. A. Purbhoo, M. Krogsgaard, M. M. Davis, Direct observation of ligand recognition by T cells. Nature 419, 845–849 (2002).

3. M. A. Purbhoo, D. J. Irvine, J. B. Huppa, M. M. Davis, T cell killing does not require the formation of a stable mature immunological synapse. Nat Immunol 5, 524–530 (2004).

4. A. Grakoui, S. K. Bromley, C. Sumen, M. M. Davis, A. S. Shaw, P. M. Allen, M. L. Dustin, The immunological synapse: a molecular machine controlling T cell activation. Science 285, 221–227 (1999).

5. Z. Ma, K. A. Sharp, P. A. Janmey, T. H. Finkel, Surface-anchored monomeric agonist pMHCs alone trigger TCR with high sensitivity. PLoS Biol. 6, e43 (2008).

6. S. J. Davis, P. A. van der Merwe, The kinetic-segregation model: TCR triggering and beyond. Nat. Immunol. 7, 803–809 (2006).

7. T. Yokosuka, K. Sakata-Sogawa, W. Kobayashi, M. Hiroshima, A. Hashimoto-Tane, M. Tokunaga, M. L. Dustin, T. Saito, Newly generated T cell receptor microclusters initiate and sustain T cell activation by recruitment of Zap70 and SLP-76. Nat. Immunol. 6, 1253–1262 (2005).

8. E. Cai, K. Marchuk, P. Beemiller, C. Beppler, M. G. Rubashkin, V. M. Weaver, A. Gérard, T.-L. Liu, B.-C. Chen, E. Betzig, F. Bartumeus, M. F. Krummel, Visualizing dynamic microvillar search and stabilization during ligand detection by T cells. Science 356, eaal3118 (2017).

9. K. H. Hu, M. J. Butte, T cell activation requires force generation. J. Cell Biol. 213, 535–542 (2016).

10. C. Hivroz, M. Saitakis, Biophysical Aspects of T Lymphocyte Activation at the Immune Synapse. Front Immunol 7, 46 (2016).

11. J. Göhring, F. Kellner, L. Schrangl, R. Platzer, E. Klotzsch, H. Stockinger, J. B. Huppa, G. J. Schütz, Temporal Analysis of T-Cell Receptor-Imposed Forces via Quantitative Single Molecule FRET Measurements. Nature Communications, doi: 10.1038/s41467-021-22775-z (2021).

12. Z. Ma, T. H. Finkel, T cell receptor triggering by force. Trends Immunol. 31, 1–6 (2010).

13. S. T. Kim, K. Takeuchi, Z.-Y. J. Sun, M. Touma, C. E. Castro, A. Fahmy, M. J. Lang, G. Wagner, E. L. Reinherz, The αβ T Cell Receptor Is an Anisotropic Mechanosensor. J. Biol. Chem. 284, 31028– 31037 (2009).

14. Y.-C. Li, B.-M. Chen, P.-C. Wu, T.-L. Cheng, L.-S. Kao, M.-H. Tao, A. Lieber, S. R. Roffler, Cutting Edge: Mechanical Forces Acting on T Cells Immobilized via the TCR Complex Can Trigger TCR Signaling. The Journal of Immunology 184, 5959–5963 (2010).

15. J. B. Huppa, M. Axmann, M. A. Mörtelmaier, B. F. Lillemeier, E. W. Newell, M. Brameshuber, L. O. Klein, G. J. Schütz, M. M. Davis, TCR-peptide-MHC interactions in situ show accelerated kinetics and increased affinity. Nature 463, 963–967 (2010).

16. B. H. Hosseini, I. Louban, D. Djandji, G. H. Wabnitz, J. Deeg, N. Bulbuc, Y. Samstag, M. Gunzer, J. P. Spatz, G. J. Hämmerling, Immune synapse formation determines interaction forces between T cells and antigen-presenting cells measured by atomic force microscopy. PNAS 106, 17852–17857 (2009).

17. J. Huang, V. I. Zarnitsyna, B. Liu, L. J. Edwards, N. Jiang, B. D. Evavold, C. Zhu, The kinetics of two-dimensional TCR and pMHC interactions determine T-cell responsiveness. Nature 464, 932–936 (2010).

18. J. Husson, K. Chemin, A. Bohineust, C. Hivroz, N. Henry, Force Generation upon T Cell Receptor Engagement. PLOS ONE 6, e19680 (2011).

19. P.-H. Puech, D. Nevoltris, P. Robert, L. Limozin, C. Boyer, P. Bongrand, Force Measurements of TCR/pMHC Recognition at T Cell Surface. PLOS ONE 6, e22344 (2011).

20. B. Liu, W. Chen, B. D. Evavold, C. Zhu, Accumulation of Dynamic Catch Bonds between TCR and Agonist Peptide-MHC Triggers T Cell Signaling. Cell 157, 357–368 (2014).

21. D. K. Das, Y. Feng, R. J. Mallis, X. Li, D. B. Keskin, R. E. Hussey, S. K. Brady, J.-H. Wang, G. Wagner, E. L. Reinherz, M. J. Lang, Force-dependent transition in the T-cell receptor β-subunit allosterically regulates peptide discrimination and pMHC bond lifetime. Proc Natl Acad Sci USA 112, 1517–1522 (2015).

22. J. Hong, S. P. Persaud, S. Horvath, P. M. Allen, B. D. Evavold, C. Zhu, Force-Regulated In Situ TCR-Peptide-Bound MHC Class II Kinetics Determine Functions of CD4+ T Cells. J. Immunol. 195, 3557–3564 (2015).

23. R. J. Mallis, K. Bai, H. Arthanari, R. E. Hussey, M. Handley, Z. Li, L. Chingozha, J. S. Duke-Cohan, H. Lu, J.-H. Wang, C. Zhu, G. Wagner, E. L. Reinherz, Pre-TCR ligand binding impacts thymocyte development before αβTCR expression. Proc Natl Acad Sci U S A 112, 8373–8378 (2015).

24. B. Liu, W. Chen, K. Natarajan, Z. Li, D. H. Margulies, C. Zhu, The cellular environment regulates in situ kinetics of T-cell receptor interaction with peptide major histocompatibility complex. Eur. J. Immunol. 45, 2099–2110 (2015).

25. L. V. Sibener, R. A. Fernandes, E. M. Kolawole, C. B. Carbone, F. Liu, D. McAffee, M. E. Birnbaum, X. Yang, L. F. Su, W. Yu, S. Dong, M. H. Gee, K. M. Jude, M. M. Davis, J. T. Groves, W. A. Goddard, J. R. Heath, B. D. Evavold, R. D. Vale, K. C. Garcia, Isolation of a Structural Mechanism for Uncoupling T Cell Receptor Signaling from Peptide-MHC Binding. Cell 174, 672–687.e27 (2018).

26. G. I. Bell, Models for the specific adhesion of cells to cells. Science 200, 618–627 (1978).

27. W. E. Thomas, E. Trintchina, M. Forero, V. Vogel, E. V. Sokurenko, Bacterial adhesion to target cells enhanced by shear force. Cell 109, 913–923 (2002).

28. B. T. Marshall, M. Long, J. W. Piper, T. Yago, R. P. McEver, C. Zhu, Direct observation of catch bonds involving cell-adhesion molecules. Nature 423, 190–193 (2003).

29. E. M. Kolawole, R. Andargachew, B. Liu, J. R. Jacobs, B. D. Evavold, 2D Kinetic Analysis of TCR and CD8 Coreceptor for LCMV GP33 Epitopes. Front. Immunol. 9 (2018).

30. Y. Liu, L. Blanchfield, V. P.-Y. Ma, R. Andargachew, K. Galior, Z. Liu, B. Evavold, K. Salaita, DNA-based nanoparticle tension sensors reveal that T-cell receptors transmit defined pN forces to their antigens for enhanced fidelity. PNAS 113, 5610–5615 (2016).

31. R. Ma, A. V. Kellner, V. P.-Y. Ma, H. Su, B. R. Deal, J. M. Brockman, K. Salaita, DNA probes that store mechanical information reveal transient piconewton forces applied by T cells. Proc Natl Acad Sci USA 116, 16949–16954 (2019).

32. V. P.-Y. Ma, Y. Liu, L. Blanchfield, H. Su, B. D. Evavold, K. Salaita, Ratiometric Tension Probes for Mapping Receptor Forces and Clustering at Intermembrane Junctions. Nano Lett. 16, 4552– 4559 (2016).

33. L. Limozin, M. Bridge, P. Bongrand, O. Dushek, P. A. van der Merwe, P. Robert, TCR–pMHC kinetics under force in a cell-free system show no intrinsic catch bond, but a minimal encounter duration before binding. Proc Natl Acad Sci USA 116, 16943–16948 (2019).

34. J. Pettmann, L. Awada, B. Różycki, A. Huhn, S. Faour, M. Kutuzov, L. Limozin, T. R. Weikl, P. A. van der Merwe, P. Robert, O. Dushek, Mechanical forces impair antigen discrimination by reducing differences in T-cell receptor/peptide–MHC off-rates. The EMBO Journal 42, e111841 (2023).

35. J. B. Huppa, G. J. Schütz, T-cell antigen recognition: catch-as-catch-can or catch-22? The EMBO Journal 42, e113507 (2023).

36. L. Schrangl, J. Göhring, F. Kellner, J. B. Huppa, G. J. Schütz, “Measurement of Forces Acting on Single T-Cell Receptors” in Imaging Cell Signaling, C. Wuelfing, R. F. Murphy, Eds. (Springer US, New York, NY, 2024; 10.1007/978-1-0716-3834-7_11), pp. 147–165.

37. M. E. Birnbaum, J. L. Mendoza, D. K. Sethi, S. Dong, J. Glanville, J. Dobbins, E. Özkan, M. M. Davis, K. W. Wucherpfennig, K. C. Garcia, Deconstructing the peptide-MHC specificity of T cell recognition. Cell 157, 1073–1087 (2014).

38. R. M. Pielak, G. P. O’Donoghue, J. J. Lin, K. N. Alfieri, N. C. Fay, S. T. Low-Nam, J. T. Groves, Early T cell receptor signals globally modulate ligand:receptor affinities during antigen discrimination. Proc. Natl. Acad. Sci. U.S.A. 114, 12190–12195 (2017).

39. A. N. Kapanidis, N. K. Lee, T. A. Laurence, S. Doose, E. Margeat, S. Weiss, Fluorescence-aided molecule sorting: Analysis of structure and interactions by alternating-laser excitation of single molecules. Proceedings of the National Academy of Sciences 101, 8936–8941 (2004).

40. L. Schrangl, J. Göhring, F. Kellner, J. B. Huppa, G. J. Schütz, Automated Two-dimensional Spatiotemporal Analysis of Mobile Single-molecule FRET Probes. JoVE (Journal of Visualized Experiments), e63124 (2021).

41. M. C. Schneider, G. J. Schütz, Don’t Be Fooled by Randomness: Valid p-Values for Single Molecule Microscopy. Front. Bioinform. 2 (2022).

42. M. Axmann, J. B. Huppa, M. M. Davis, G. J. Schütz, Determination of Interaction Kinetics between the T Cell Receptor and Peptide-Loaded MHC Class II via Single-Molecule Diffusion Measurements. Biophysical Journal 103, L17–L19 (2012).

43. G. P. O’Donoghue, R. M. Pielak, A. A. Smoligovets, J. J. Lin, J. T. Groves, Direct single molecule measurement of TCR triggering by agonist pMHC in living primary T cells. Elife 2, e00778 (2013).

44. L. Schrangl, V. Mühlgrabner, R. Platzer, F. Kellner, J. Wieland, R. Obst, J. L. Toca-Herrera, J. B. Huppa, G. J. Schütz, J. Göhring, Advanced Quantification of Receptor–Ligand Interaction Lifetimes via Single-Molecule FRET Microscopy. Biomolecules 14, 1001 (2024).

45. J. Göhring, L. Schrangl, G. J. Schütz, J. B. Huppa, Mechanosurveillance: Tiptoeing T Cells. Frontiers in Immunology 13 (2022).

46. O. Acuto, T-cell virtuosity in “knowing thyself”. Front Immunol 15, 1343575 (2024).

47. M. Fritzsche, K. Kruse, Mechanical force matters in early T cell activation. Proc. Natl. Acad. Sci. U.S.A. 121, e2404748121 (2024).

48. Y. Hu, J. Rogers, Y. Duan, A. Velusamy, S. Narum, S. Al Abdullatif, K. Salaita, Quantifying T cell receptor mechanics at membrane junctions using DNA origami tension sensors. Nat. Nanotechnol., 1–12 (2024).

49. O. Milstein, S.-Y. Tseng, T. Starr, J. Llodra, A. Nans, M. Liu, M. K. Wild, P. A. van der Merwe, D. L. Stokes, Y. Reisner, M. L. Dustin, Nanoscale Increases in CD2-CD48-mediated Intermembrane Spacing Decrease Adhesion and Reorganize the Immunological Synapse. J. Biol. Chem. 283, 34414–34422 (2008).

50. M. Fölser, V. Motsch, R. Platzer, J. B. Huppa, G. J. Schütz, A Multimodal Platform for Simultaneous T-cell Imaging, Defined Activation, and Mechanobiological Characterization. Cells 10, 235 (2021).

51. J. B. Huppa, M. Gleimer, C. Sumen, M. M. Davis, Continuous T cell receptor signaling required for synapse maintenance and full effector potential. Nat. Immunol. 4, 749–755 (2003).

52. R. Platzer, J. Hellmeier, J. Göhring, I. D. Perez, P. Schatzlmaier, C. Bodner, M. Focke-Tejkl, G. J. Schütz, E. Sevcsik, H. Stockinger, M. Brameshuber, J. B. Huppa, Monomeric agonist peptide/MHCII complexes activate T-cells in an autonomous fashion. EMBO Rep 24, e57842 (2023).

53. M. Brameshuber, F. Kellner, B. K. Rossboth, H. Ta, K. Alge, E. Sevcsik, J. Göhring, M. Axmann, F. Baumgart, N. R. J. Gascoigne, S. J. Davis, H. Stockinger, G. J. Schütz, J. B. Huppa, Monomeric TCRs drive T cell antigen recognition. Nat. Immunol. 19, 487–496 (2018).

54. M. Howarth, D. J.-F. Chinnapen, K. Gerrow, P. C. Dorrestein, M. R. Grandy, N. L. Kelleher, A. El-Husseini, A. Y. Ting, A monovalent streptavidin with a single femtomolar biotin binding site. Nat Methods 3, 267–273 (2006).

55. L. Schrangl, Single-molecule FRET analysis software (v3.0), version 3.0, Zenodo (2021); 10.5281/zenodo.5115967.

56. P. Virtanen, R. Gommers, T. E. Oliphant, M. Haberland, T. Reddy, D. Cournapeau, E. Burovski, P. Peterson, W. Weckesser, J. Bright, S. J. van der Walt, M. Brett, J. Wilson, K. J. Millman, N. Mayorov, A. R. J. Nelson, E. Jones, R. Kern, E. Larson, C. J. Carey, İ. Polat, Y. Feng, E. W. Moore, J. VanderPlas, D. Laxalde, J. Perktold, R. Cimrman, I. Henriksen, E. A. Quintero, C. R. Harris, A. M. Archibald, A. H. Ribeiro, F. Pedregosa, P. van Mulbregt, SciPy 1.0 Contributors, SciPy 1.0: fundamental algorithms for scientific computing in Python. Nat Methods 17, 261–272 (2020).

57. L. Schrangl, sdt-python: Python library for fluorescence microscopy data anlysis, version 16.1, Zenodo (2021); 10.5281/zenodo.4604494.

58. L. Schrangl, smfret-bondtime: Quantification of receptor–ligand interaction times via single-molecule FRET, version 1.0.1, Zenodo (2024); 10.5281/zenodo.13255170.

59. R. Mw, L. Jj, H. B, Assessment of Fura-2 for measurements of cytosolic free calcium. Cell calcium 11 (1990).

