## Supplementary Materials for "CD4+ T-cells create a stable mechanical environment for force-sensitive TCR:pMHC interactions"

#### Fig. S1

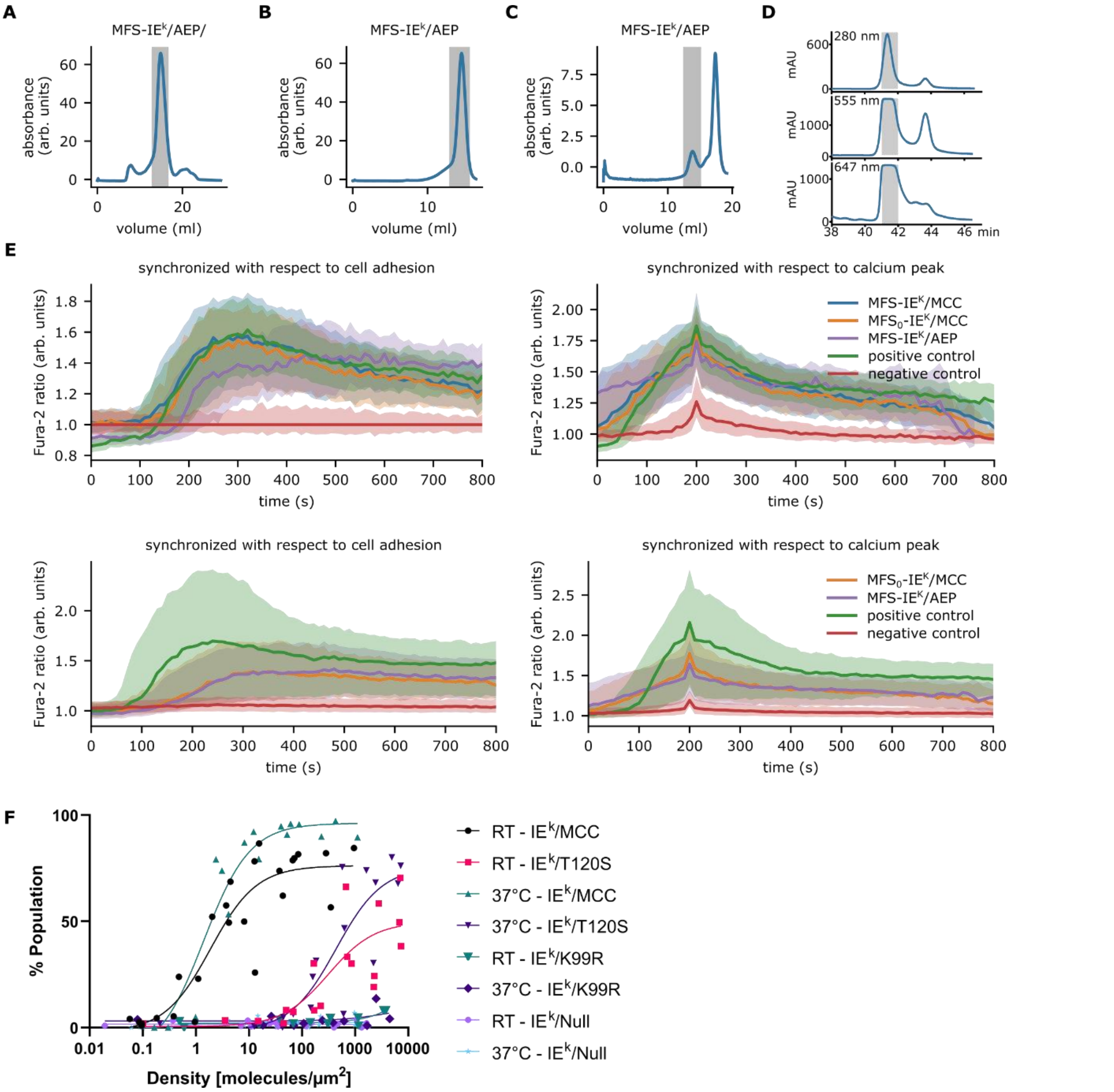


**Fig. S1. Synthesis and functional validation of the AEP-IEk molecular force sensor. (A)** Refolded murine pMHC class II molecule IEk with AEP peptide ($IE^k$/AEP) was purified by Superdex 200 10/300 GL (S200) size exclusion chromatography. The grey areas correspond to collected fractions. **(B)** Post conjugation, the surplus of DBCO was removed from monomeric $IE^k$/AEP-DBCO via Superdex 75 10/300 GL (S75) gel filtration. **(C)** Unconjugated MFS was separated from MFS-$IE^k$/AEP via S75 gel filtration. **(D)** Absorbance profiles of HPLC purified Alexa 555 /Alexa 647 dual-labelled AEP peptide confirmed successful label conjugation. **(E)** Calcium flux analysis of the SLBs decorated with MFS-$IE^k$/MCC, MFS-$IE^k$/AEP, and unlabeled $MFS_0$-$IE^k$/MCC, as well as ICAM-1 and B7-1. Positive control: ICAM-1, B7-1 and his-tagged $IE^k$/MCC. Negative control: only ICAM-1 and B7-1. Two independent experiments performed at room temperature are shown (upper and lower panel). **(F)** Measurement of T-cell sensitivity in room temperature vs. 37°C using his-tagged IEk with the strong agonistic MCC, weak agonistic T102S, antagonistic K99R, and Null peptide. Calcium response was measured for varying antigen densities. For comparability, data sets for varying temperatures were measured on the same day and data was pooled from three independent experiments.

Fig. S2

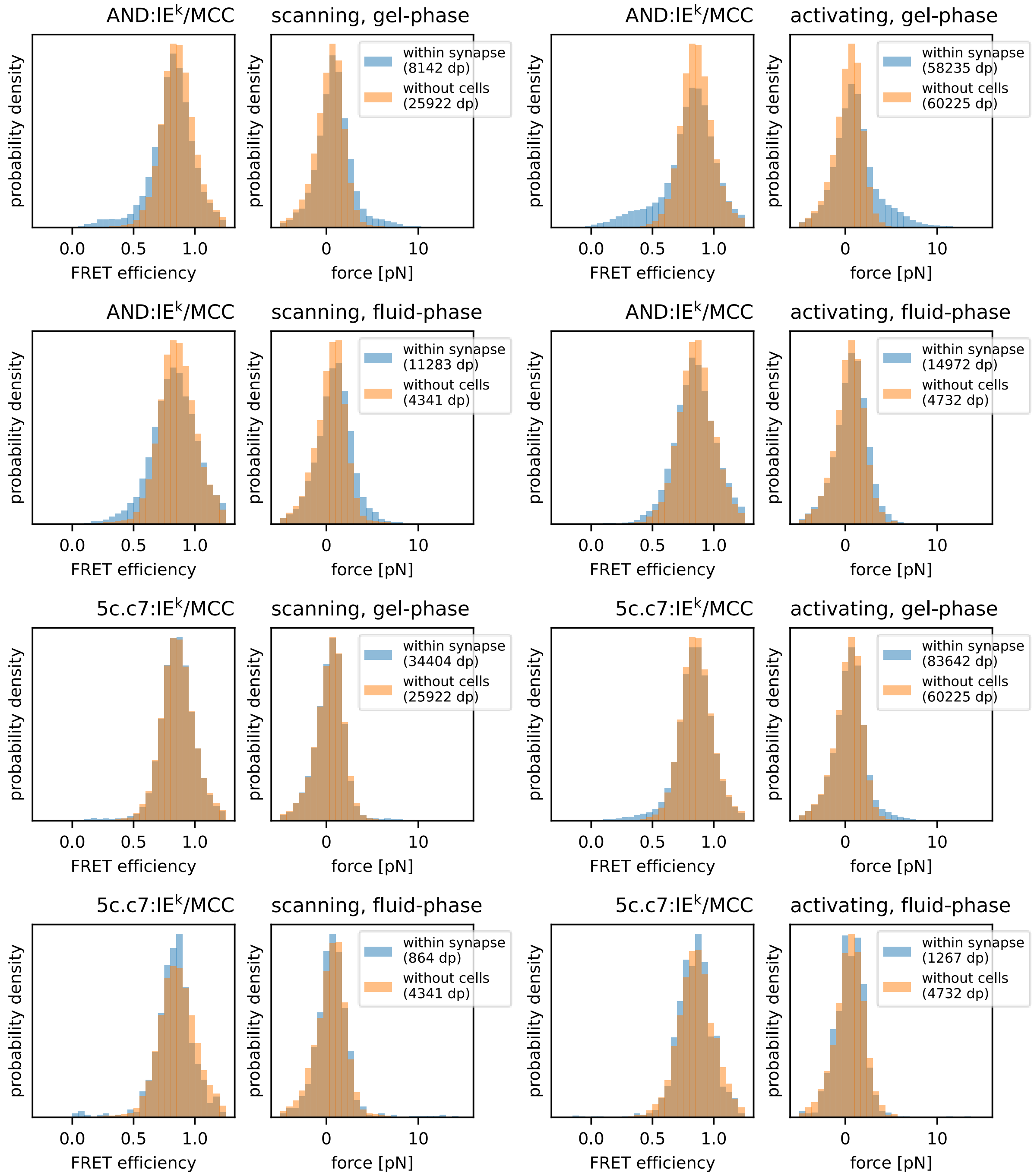
AND:IEk/MCC
scanning, gel-phase
activating, gel-phase
scanning, fluid-phase
activating, fluid-phase
5c.c7:IEk/MCC
probability density
FRET efficiency
force [pN]
within synapse
(8142 dp)
without cells
(25922 dp)
(58235 dp)
(60225 dp)
(11283 dp)
(4341 dp)
(14972 dp)
(4732 dp)
(34404 dp)
(83642 dp)
(864 dp)
(1267 dp)

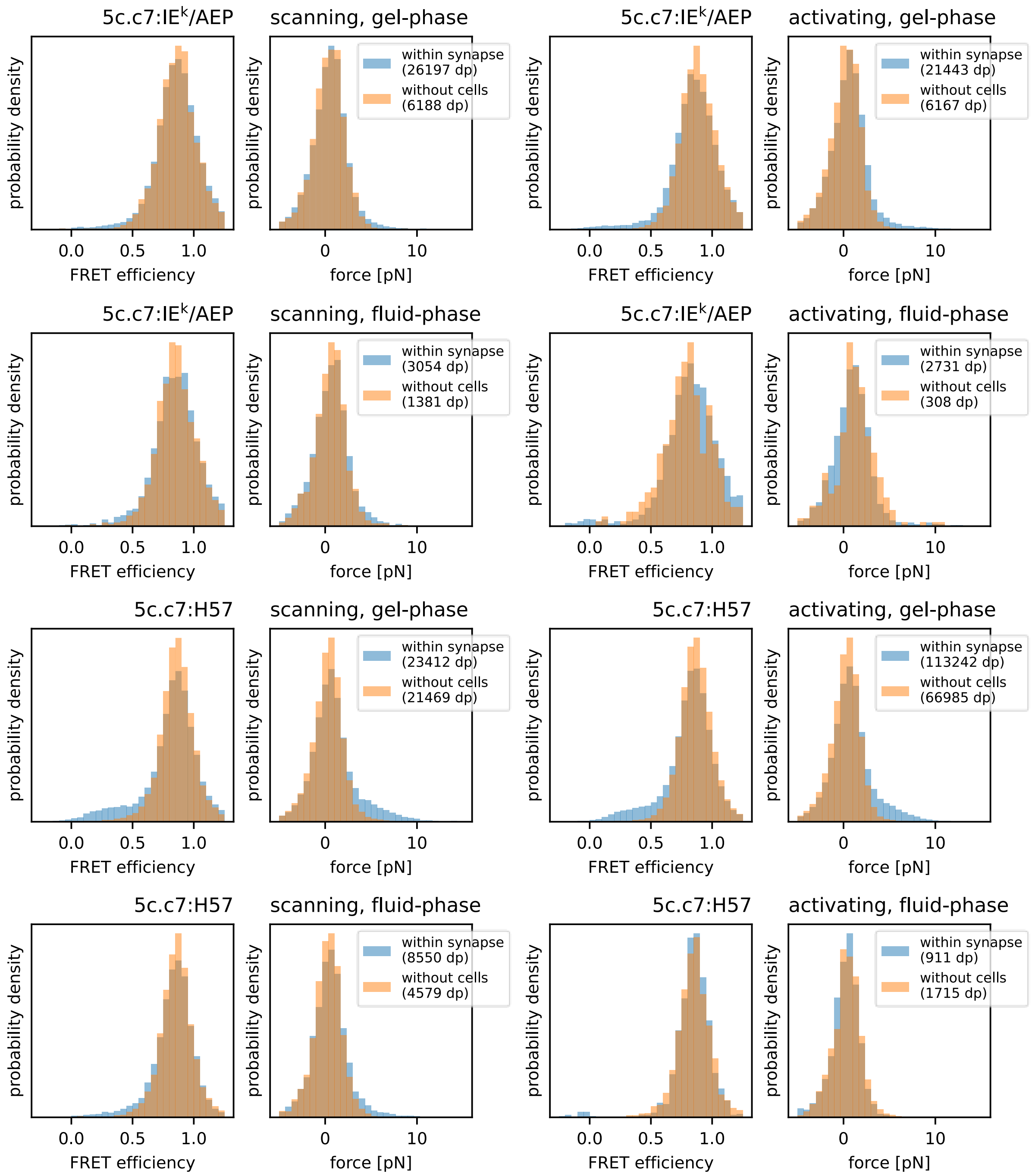


**Fig. S2. FRET efficiency and force histograms of the single molecule FRET trajectories for scanning (left side of the panel) and activating conditions (right side of the panel) on gel-phase and fluid SLBs.** The FRET efficiency histograms are plotted in the first and third column, whereas the respective force histograms are plotted in the second and fourth column. The number of data points participating in the analysis are mentioned in the legend, and the number of trajectories extracted from independent experiments are summarized in Table S1.

Fig. S3

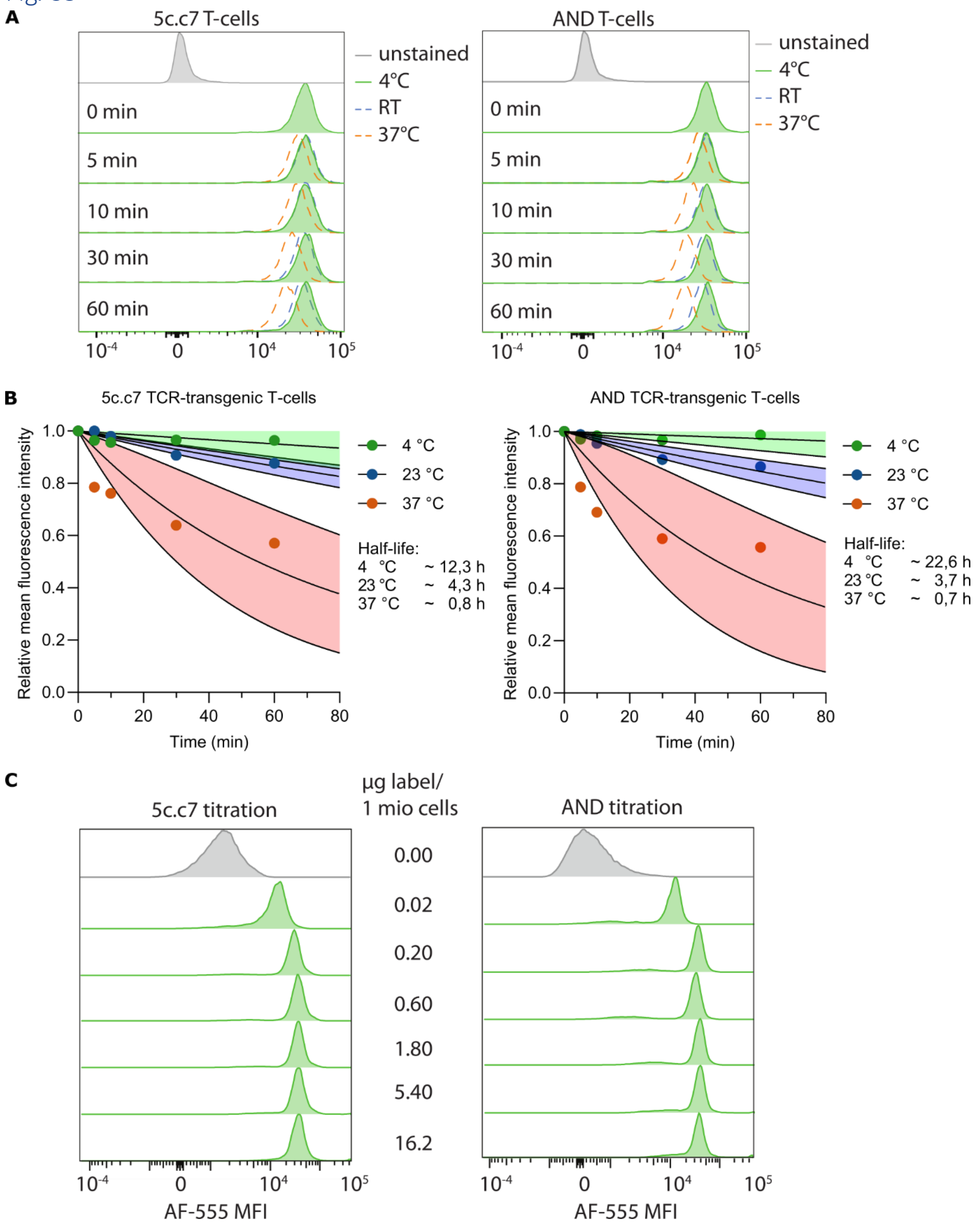


**Fig. S3. Flow cytometry data for the label stability and saturation of transgenic T-cells with fluorescent H57-scF$_V$. (A)** 5c.c7 and AND T-cells were labelled and incubated for up to 1 hours in 4°C, 24°C and 37°C. Label density was detected via flow cytometry. **(B)** Halftime estimation of bound label for data in panel A. Data points were fitted with a one phase decay curve with a plateau constraint of 0.1. 95% confidence intervals of the fit are shown. Each histogram represents approximately 50000 cells from one experiment. **(C)** Label titration to achieve saturation using 5c.c7 and AND T-cells.

Fig. *S4*

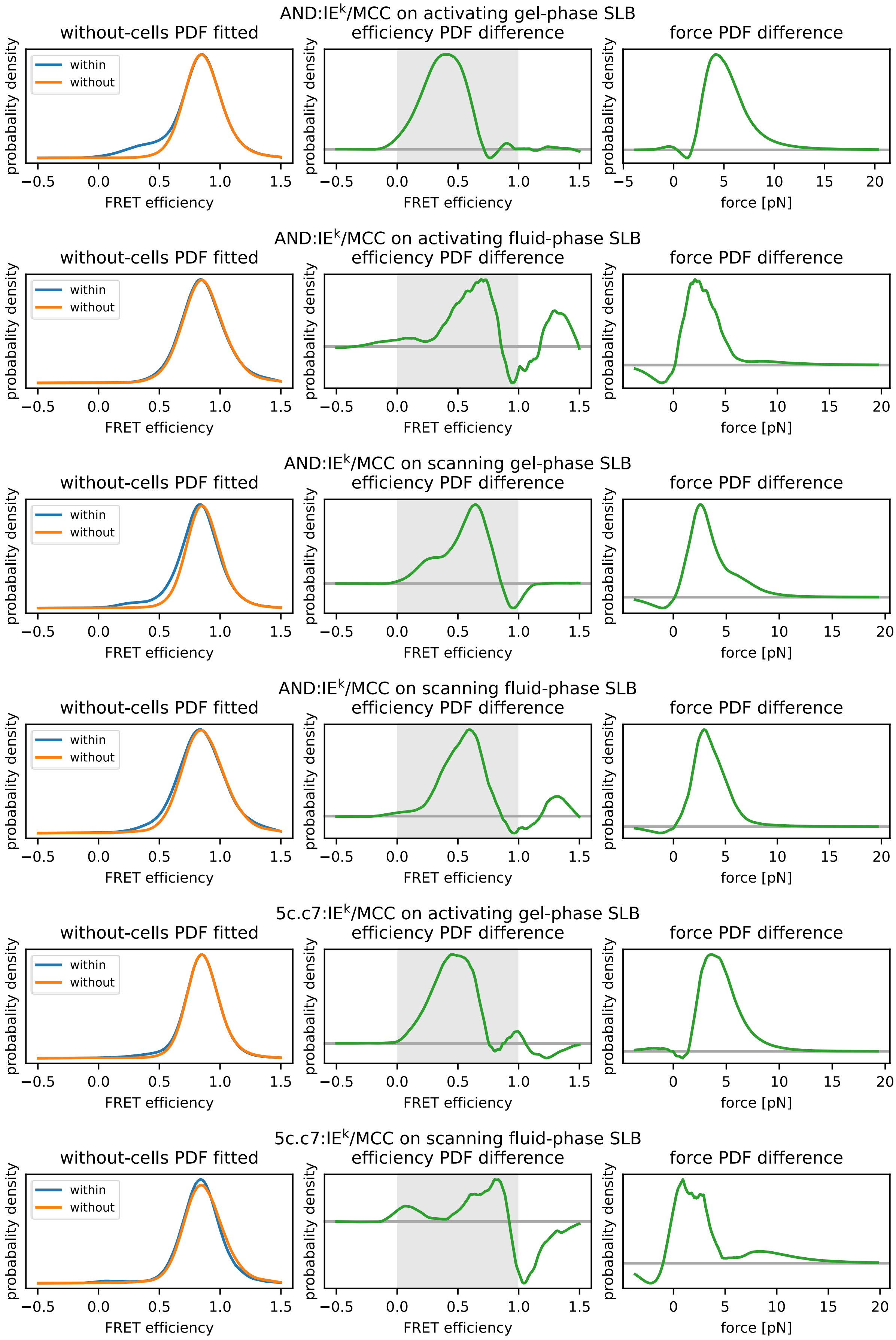
AND:IE$^k$/MCC on activating gel-phase SLB
AND:IE$^k$/MCC on activating fluid-phase SLB
AND:IE$^k$/MCC on scanning gel-phase SLB
AND:IE$^k$/MCC on scanning fluid-phase SLB
5c.c7:IE$^k$/MCC on activating gel-phase SLB
5c.c7:IE$^k$/MCC on scanning fluid-phase SLB
without-cells PDF fitted
efficiency PDF difference
force PDF difference
within
without
probabality density
FRET efficiency
force [pN]

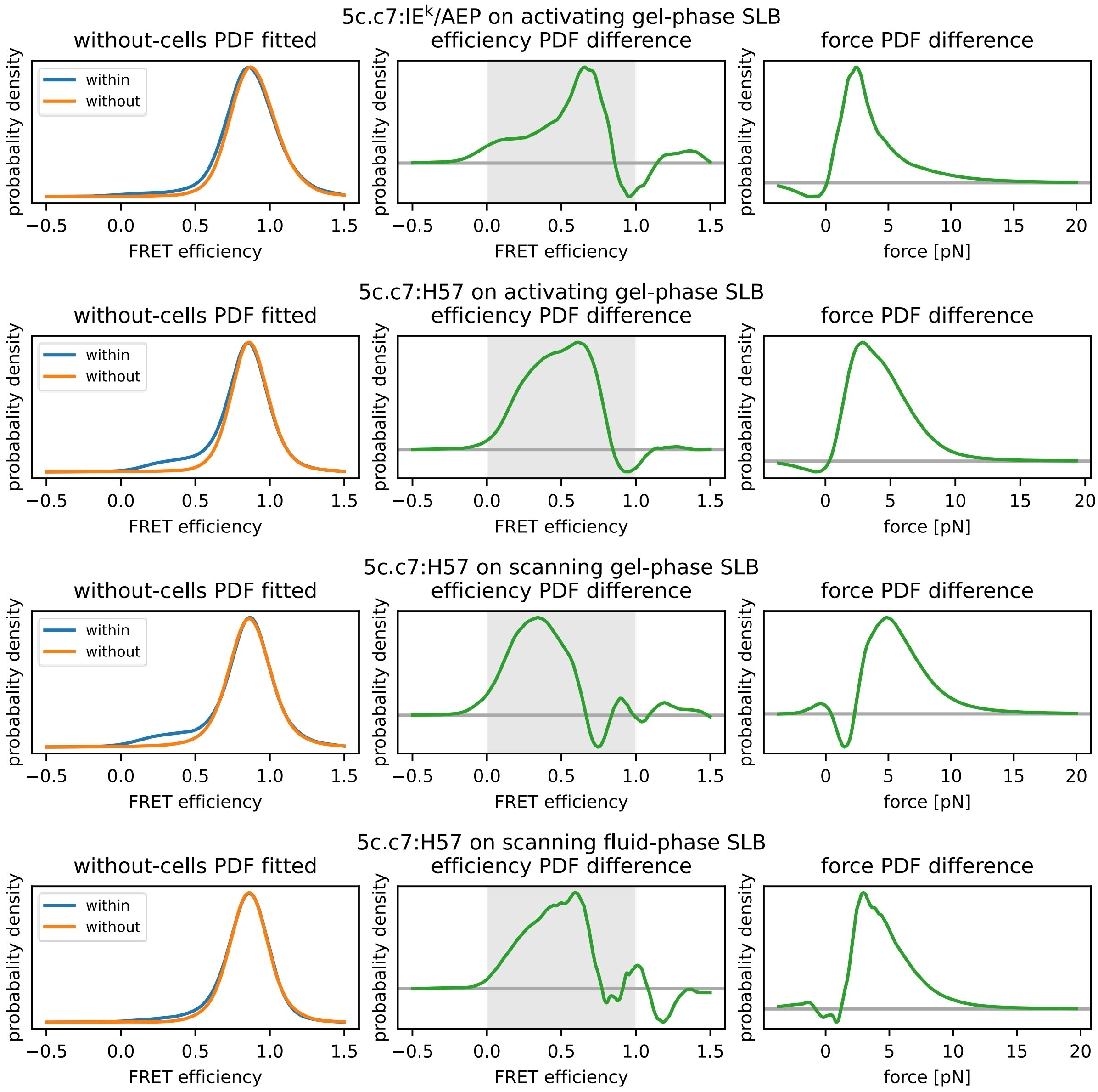


**Fig. S4. Extraction of the low frequency force events from the entire population of FRET trajectories via PDF subtraction of the no-cell data. (left panel)** FRET efficiency histograms of raw data. **(middle panel)** Result of the PDF subtraction. **(right panel)** Conversion of the low FRET population into force. The number of trajectories extracted from independent experiments are summarized in Table S1.

Fig. S5

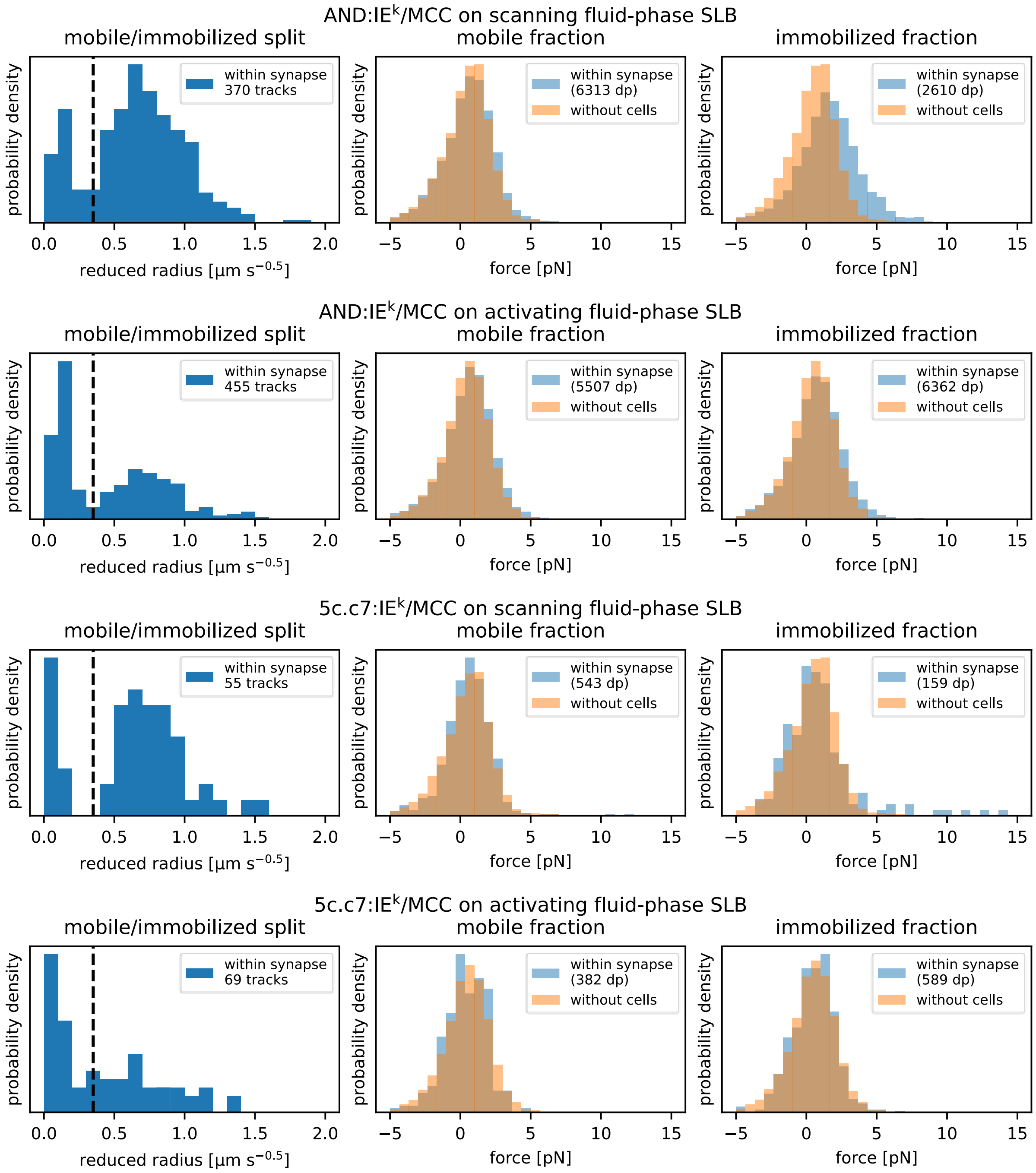
AND:IE$^k$/MCC on scanning fluid-phase SLB
mobile/immobilized split
mobile fraction
immobilized fraction
within synapse 370 tracks
within synapse (6313 dp)
without cells
within synapse (2610 dp)
without cells
AND:IE$^k$/MCC on activating fluid-phase SLB
within synapse 455 tracks
within synapse (5507 dp)
within synapse (6362 dp)
5c.c7:IE$^k$/MCC on scanning fluid-phase SLB
within synapse 55 tracks
within synapse (543 dp)
within synapse (159 dp)
5c.c7:IE$^k$/MCC on activating fluid-phase SLB
within synapse 69 tracks
within synapse (382 dp)
within synapse (589 dp)
probability density
reduced radius [μm s$^{-0.5}$]
force [pN]

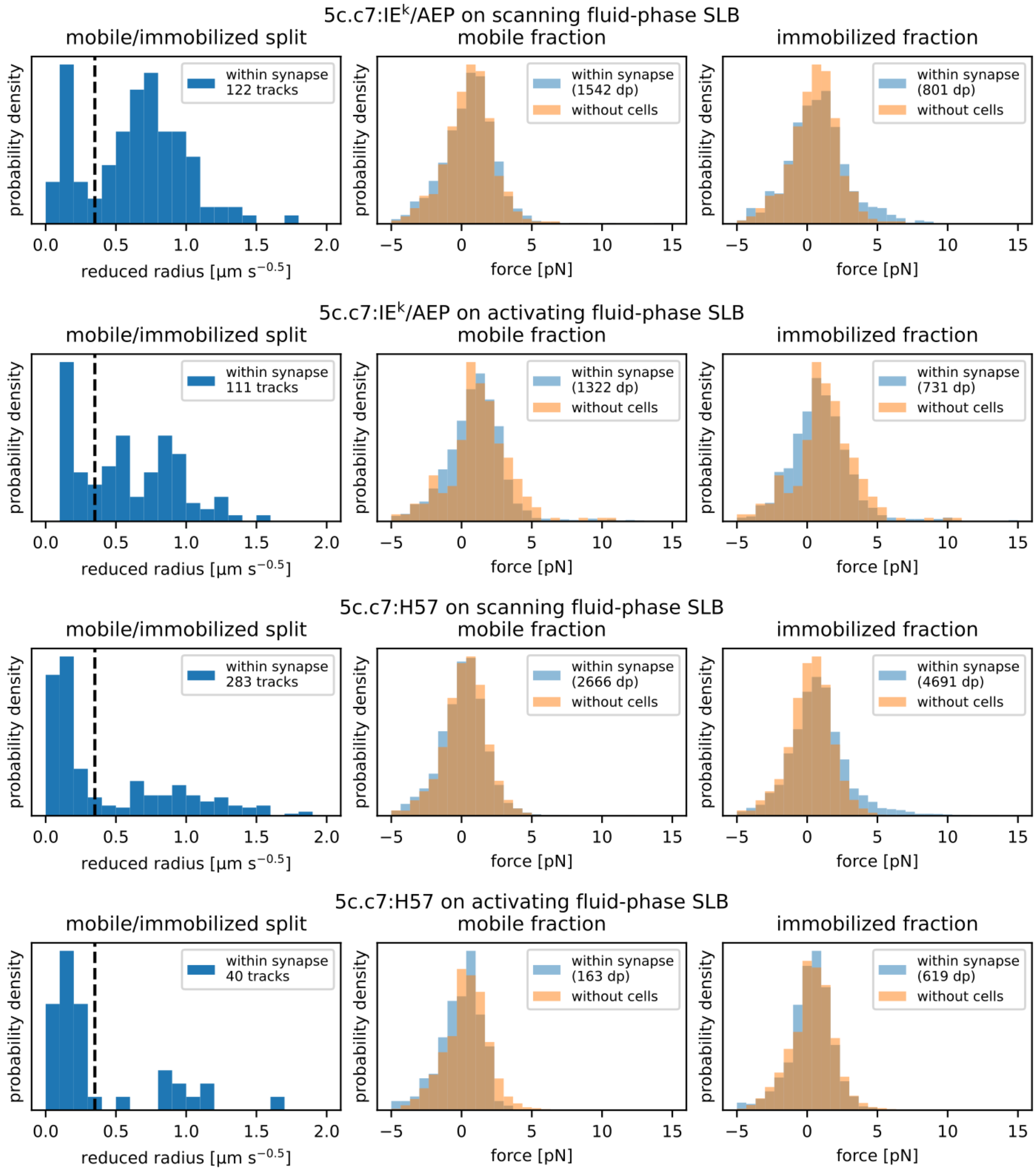


***Fig. S5.*** **Force histograms of the mobile and immobilized fraction and the respective immobilizations plots.** The number of trajectories and data points participating in the analysis are mentioned in the legend.

Fig. S6

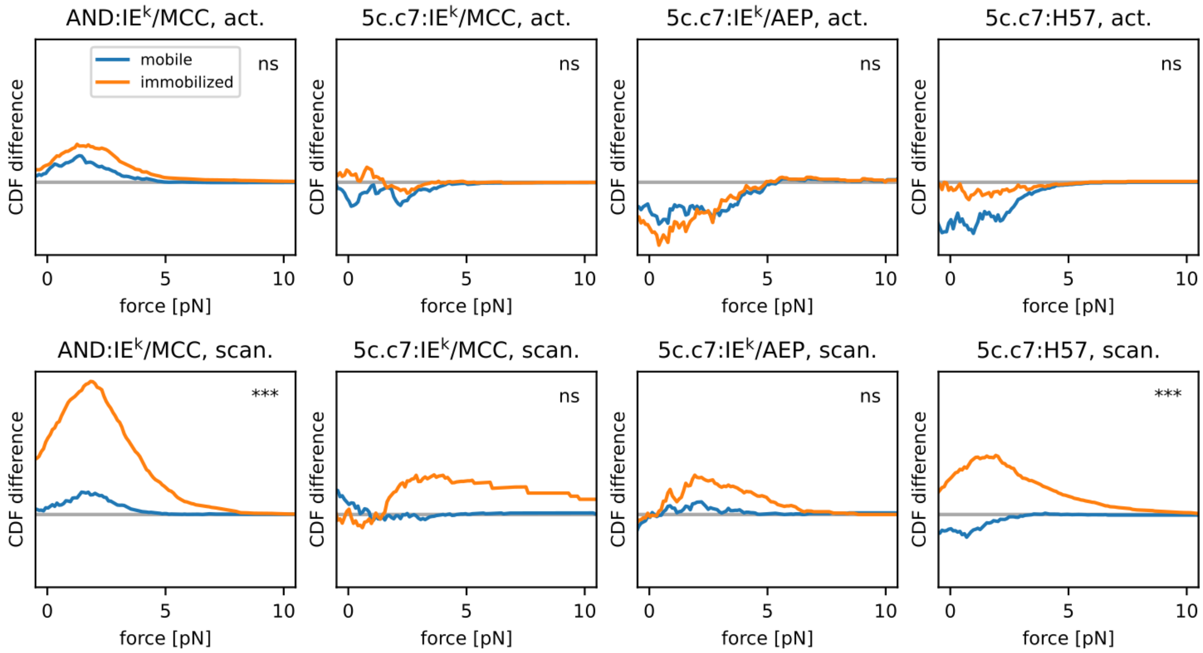


***Fig. S6.*** Comparison of the cumulative density function (CDF) of the mobile and immobilized fraction (no-cell background was subtracted). Significant differences to the no-cell data are indicated by the asterisks in the corner of each plot. * - p-value < 0.05; ** - p-value < 0.01; *** - p-value < 0.001; ns – not significant.

#### Table S1

| | | counts | | | high-force fraction | | | high-force quartiles [pN] | | |
|---|---|---|---|---|---|---|---|---|---|---|
| | p-value | data points | tracks | videos | threshold | proportion | error | Q1 | Q2 | Q3 |
| 5c.c7:IE$^k$/AEP, activating gel-phase SLB | 0.0006 | 21443 | 1091 | 339 | 0.65 | 0.06 | 0.01 | 2.0 | 3.0 | 4.9 |
| 5c.c7:IE$^k$/AEP, activating fluid-phase SLB | 0.8113 | 2731 | 134 | 79 | N/A | N/A | N/A | N/A | N/A | N/A |
| 5c.c7:IE$^k$/AEP, scanning gel-phase SLB | 0.3188 | 26197 | 1116 | 334 | N/A | N/A | N/A | N/A | N/A | N/A |
| 5c.c7:IE$^k$/AEP, scanning fluid-phase SLB | 0.3473 | 3054 | 143 | 111 | N/A | N/A | N/A | N/A | N/A | N/A |
| 5c.c7:H57, activating gel-phase SLB | 0.0001 | 113242 | 3879 | 1626 | 0.65 | 0.11 | 0.01 | 2.7 | 4.0 | 5.7 |
| 5c.c7:H57, activating fluid-phase SLB | 0.6267 | 911 | 49 | 44 | N/A | N/A | N/A | N/A | N/A | N/A |
| 5c.c7:H57, scanning gel-phase SLB | 0.0001 | 23412 | 1029 | 543 | 0.63 | 0.11 | 0.01 | 4.2 | 5.5 | 7.2 |
| 5c.c7:H57, scanning fluid-phase SLB | 0.0175 | 8550 | 320 | 226 | 0.64 | 0.05 | 0.02 | 3.0 | 4.2 | 5.9 |
| 5c.c7:IE$^k$/MCC, activating gel-phase SLB | 0.0001 | 83642 | 2746 | 1265 | 0.65 | 0.04 | 0.00 | 3.2 | 4.2 | 5.5 |
| 5c.c7:IE$^k$/MCC, activating fluid-phase SLB | 0.0517 | 1267 | 113 | 98 | N/A | N/A | N/A | N/A | N/A | N/A |
| 5c.c7:IE$^k$/MCC, scanning gel-phase SLB | 0.3914 | 34404 | 1054 | 553 | N/A | N/A | N/A | N/A | N/A | N/A |
| 5c.c7:IE$^k$/MCC, scanning fluid-phase SLB | 0.0029 | 864 | 75 | 66 | 0.62 | 0.02 | 0.02 | 0.9 | 2.2 | 3.9 |
| AND:IE$^k$/MCC, activating gel-phase SLB | 0.0001 | 58235 | 1714 | 529 | 0.65 | 0.14 | 0.01 | 3.8 | 5.0 | 6.5 |
| AND:IE$^k$/MCC, activating fluid-phase SLB | 0.0137 | 14972 | 482 | 289 | 0.62 | 0.04 | 0.01 | 1.8 | 2.8 | 4.0 |
| AND:IE$^k$/MCC, scanning gel-phase SLB | 0.0001 | 8142 | 292 | 185 | 0.64 | 0.08 | 0.02 | 2.2 | 3.1 | 4.6 |
| AND:IE$^k$/MCC, scanning fluid-phase SLB | 0.0003 | 11283 | 409 | 239 | 0.62 | 0.06 | 0.02 | 2.5 | 3.4 | 4.5 |

**Table S1: Summary of statistics for all recorded molecular force data sets.**

#### Table S2

| | SLB | T [°C] | lifetime [s] estimate | lifetime [s] error | recording intervals [s] | counts by interval: frames per movie | counts by interval: recorded tracks |
|---|---|---|---|---|---|---|---|
| 5c.c7:IE$^k$/AEP | | | | | | | |
| 0 | gel-phase | 23 | 152.2 | 27.0 | 1.0, 2.5, 5.0, 7.5 | 180, 75, 40, 35 | 148, 162, 80, 161 |
| 0 | fluid-phase | 23 | 191.8 | 78.5 | 1.0, 2.5, 5.0, 7.5 | 180, 75, 40, 35 | 74, 105, 91, 75 |
| 1 | gel-phase | 23 | 102.2 | 15.2 | 1.0, 2.5, 5.0, 7.5 | 180, 75, 40, 35 | 157, 175, 219, 218 |
| 1 | fluid-phase | 23 | 88.9 | 12.1 | 1.0, 2.5, 5.0, 7.5 | 180, 75, 40, 35 | 151, 97, 185, 158 |
| 5c.c7:IE$^k$/MCC | | | | | | | |
| 2 | gel-phase | 27 | 4.5 | 0.8 | 0.05, 0.1, 0.25, 0.5, 1.0 | 1000, 500, 200, 100, 50 | 32, 51, 49, 54, 50 |
| 2 | fluid-phase | 27 | 4.1 | 0.5 | 0.05, 0.1, 0.25, 0.5 | 1000, 500, 200, 100 | 160, 149, 125, 89 |
| 3 | gel-phase | 27 | 4.3 | 0.8 | 0.05, 0.175, 0.5, 1.5 | 500, 250, 100, 50 | 19, 24, 39, 43 |
| 3 | fluid-phase | 27 | 5.3 | 0.8 | 0.05, 0.175, 0.5, 1.5 | 500, 250, 100, 50 | 38, 42, 56, 52 |
| 4 | gel-phase | 27 | 4.2 | 1.0 | 0.05, 0.1, 0.25, 0.5, 1.0, 1.5 | 500, 250, 150, 75, 25, 16 | 12, 9, 21, 35, 15, 11 |
| 4 | fluid-phase | 27 | 4.3 | 0.4 | 0.05, 0.1, 0.25, 0.5, 1.0, 1.5 | 500, 250, 150, 75, 30, 20 | 117, 138, 133, 69, 90, 67 |
| 5 | gel-phase | 23 | 18.5 | 2.1 | 0.1, 0.25, 0.5, 1.0, 2.0 | 500, 200, 100, 50, 30 | 149, 254, 181, 188, 147 |
| 5 | fluid-phase | 23 | 13.5 | 1.0 | 0.1, 0.25, 0.5, 1.0, 2.0 | 500, 200, 100, 50, 30 | 256, 408, 254, 316, 320 |
| 6 | gel-phase | 23 | 11.6 | 1.5 | 0.1, 0.25, 0.5, 1.0, 2.0 | 500, 200, 100, 50, 30 | 149, 107, 103, 88, 89 |
| 6 | fluid-phase | 23 | 10.1 | 0.7 | 0.1, 0.25, 0.5, 1.0, 2.0 | 500, 200, 100, 50, 30 | 344, 327, 360, 327, 213 |
| 7 | gel-phase | 23 | 7.0 | 1.1 | 0.1, 0.25, 0.5, 1.0, 2.0 | 500, 200, 100, 50, 30 | 75, 46, 103, 37, 33 |
| 7 | fluid-phase | 23 | 13.6 | 1.3 | 0.1, 0.25, 0.5, 1.0, 2.0 | 500, 200, 100, 50, 30 | 191, 211, 203, 180, 188 |
| AND:IE$^k$/MCC | | | | | | | |
| 8 | gel-phase | 27 | 28.5 | 5.5 | 0.25, 1.0, 2.0, 4.0, 8.0 | 300, 75, 50, 25, 15 | 42, 43, 29, 3, 19 |
| 8 | fluid-phase | 27 | 27.1 | 6.6 | 0.25, 1.0, 2.0, 4.0, 8.0 | 300, 75, 50, 25, 15 | 25, 16, 11, 14, 6 |
| 9 | gel-phase | 27 | 19.0 | 3.7 | 0.25, 1.0, 2.0, 4.0 | 300, 75, 50, 25 | 25, 14, 19, 31 |
| 9 | fluid-phase | 27 | 21.9 | 5.2 | 0.25, 1.0, 2.0, 4.0, 8.0 | 300, 75, 50, 25, 15 | 21, 14, 15, 13, 8 |
| 10 | gel-phase | 27 | 29.0 | 3.1 | 0.25, 0.5, 1.0, 2.0, 4.0 | 400, 200, 100, 75, 35 | 140, 112, 94, 153, 82 |
| 10 | fluid-phase | 27 | 35.4 | 7.0 | 0.25, 1.0, 2.0, 3.0, 4.0 | 300, 75, 75, 50, 35 | 63, 40, 24, 17, 16 |
| 11 | gel-phase | 23 | 75.6 | 12.3 | 0.25, 0.5, 1.0, 2.5, 5.0 | 500, 250, 180, 75, 40 | 80, 97, 87, 114, 70 |
| 11 | fluid-phase | 23 | 89.7 | 24.4 | 0.25, 0.5, 1.0, 2.5, 5.0 | 500, 250, 180, 75, 40 | 59, 77, 43, 64, 28 |
| 12 | gel-phase | 23 | 88.3 | 15.6 | 0.25, 0.5, 1.0, 2.5, 5.0 | 500, 250, 180, 75, 40 | 155, 146, 142, 170, 143 |
| 12 | fluid-phase | 23 | 69.7 | 14.5 | 0.25, 0.5, 1.0, 2.5, 5.0 | 500, 250, 180, 75, 40 | 122, 180, 170, 150, 73 |

**Table S2: Summary of statistics for all recorded lifetime data sets.** The experimental set is indicated by the number in the first column.

### Table S3

| | drug | T [°C] | lifetime [s] estimate | lifetime [s] error | recording intervals [s] | counts by interval: frames per movie | counts by interval: recorded tracks |
|---|---|---|---|---|---|---|---|
| 0 | Cyto-D | 23 | 11.1 | 1.8 | 0.1, 0.25, 0.5, 1.0, 2.0 | 500, 200, 100, 50, 30 | 142, 22, 34, 13, 73 |
| 1 | Cyto-D | 23 | 12.5 | 2.0 | 0.1, 0.25, 0.5, 1.0, 2.0 | 500, 200, 100, 50, 30 | 104, 99, 53, 60, 72 |
| 1 | DMSO | 23 | 9.9 | 1.7 | 0.1, 0.25, 0.5, 1.0, 2.0 | 500, 200, 100, 50, 30 | 72, 136, 153, 100, 59 |

**Table S3: Summary of statistics for recorded lifetime data sets upon actin cytoskeleton disruption.** The experimental set is indicated by the number in the first column.
